# “Broad-Spectrum Heavily Mutated Monoclonal Antibody Isolated from COVID-19 Convalescent Vaccinee with Capacity to Neutralize SARS-CoV2 Variants Ranging from B.1 to BQ.1.1.”

**DOI:** 10.1101/2023.05.04.539267

**Authors:** Alok Choudhary, David Calianese, William Honnen, Afsal Kolloli, Ryan J. Dikdan, Dabbu Jaijyan, Ge Song, Tazio Capozzola, Chandler Sy, Vikash Akkaraju, Arun Mattappallil, Anthony Rosania, Salman Khan, Mark Lerman, Afzal Nikaein, Selvakumar Subbian, Ting-Chao Chou, Raiees Andrabi, Dennis Burton, Abraham Pinter

**Affiliations:** Public Health Research Institute, Rutgers University, Newark, NJ, USA; Rutgers New Jersey Medical School, Newark, NJ, USA; Department of Immunology and Microbiology, Scripps Research, La Jolla, CA, USA; University Hospital, Newark; Dallas Nephrology Associates, Dallas, TX, USA; Medical City Specialty, Dallas, TX, USA; PD Science LLC, USA

**Keywords:** COVID-19, SARS-CoV2, Omicron subvariants, Anti-RBD, Anti-RBM, Spike, broadly neutralizing antibody, Cross-CoV2 neutralization, Hypermutated Ab, Synergism

## Abstract

Graphical Abstract

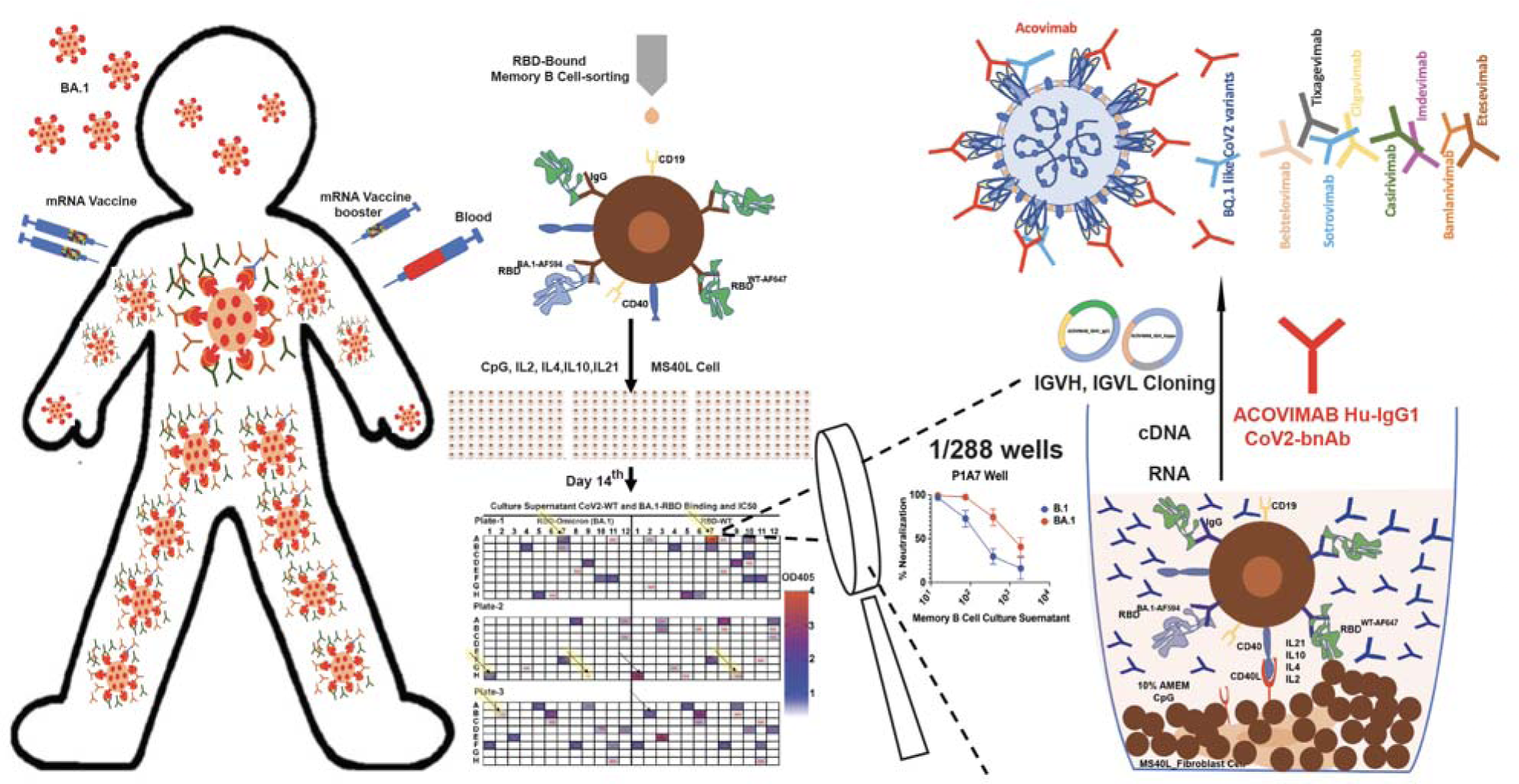

**In brief:** Choudhary et al. have isolated and characterized Acovimab, a broadly neutralizing RBM-specific human monoclonal antibody with a relatively high level of somatic hypermutation, which potently neutralizes SARS-CoV2 variants ranging from Wuhan_B.1_ to Omicron_BQ.1.1_, but not the XBB.1.5 variant. Acovimab also possesses strong synergistic neutralizing activity against some Omicron variants when combined with Sotrovimab. Polyclonal plasma antibodies from COVID-19 vaccinees who had recovered from SARS-CoV2 infection were shown to possess low neutralizing titers of antibodies against conserved RBD targets of CoV2 variants including XBB.1.5, which also synergistically neutralize with Sotrovimab against this variant.

**Summary:** The increasing prevalence of the highly antibody-evasive Omicron sublineages increases the risk of breakthrough infections and leaves high-risk and vulnerable immunocompromised individuals with no effective options for prophylactic or therapeutic antibody treatments. Here, we report a heavily mutated anti-RBD monoclonal antibody, Acovimab, directed against a site in the receptor-binding motif (RBM) region of the CoV2 receptor-binding domain (RBD), that possesses very broad and highly potent neutralizing activity against CoV2 variants, including many Omicron variants. This antibody is derived from the IGHV1-58*01 germline sequence and possesses a relatively high level of mutation (15.5% of the V_H_ aa sequence), which is unusual for anti-RBD antibodies. Neutralizing activity was very potent (IC_50s_ range of 1-9 ng/ml) for early Omicron subvariants that possess an unmutated F486 residue and is retained but less potent (IC_50s_ of 200-650 ng/ml) for more resistant Omicron subvariants which contain the F486V mutation (BA4/5, BA4.6, and BQ1.1), but is lost for the later ultra-resistant variants that contain F486S (XBB) or F486P (XBB.1.5) mutations. Based on these specificities, it is predicted that Acovimab by itself should protect against CoV2 infections other than those caused by the XBB/XBB.1.5 family. Acovimab also shows strong synergy in neutralization when combined with Sotrovimab, which neutralizes all Omicron variants, including XBB.1.5. Plasma from subjects with hybrid immunity (induced by vaccination + infection) possessed low levels of XBB.1.5 RBM-targeting plasma-neutralizing antibodies, and these also neutralized synergistically when combined with Sotrovimab. These results suggest potentially novel immunotherapeutic options for treating most of the CoV2 variants responsible for current infections.

## Highlights

- Acovimab, a SARS-CoV2 broadly neutralizing mAb, is effective in neutralizing SARS-CoV2 isolates, including variants ranging from B.1 to BQ.1.1 SARS-CoV2.
- Acovimab is heavily mutated, with 15.5% IGHV and 4.5% IGKV amino acid mutations compared to the original germline sequences.
- Acovimab targets an epitope in the RBM and synergistically neutralizes the BQ.1.1 Omicron subvariant when combined with Sotrovimab.
- Polyclonal plasma antibodies from CoV2-infected vaccines possess low titers of highly conserved RBM-specific antibodies that extend to the XBB.1.5 variant, and also possess neutralization synergy when combined with Sotrovimab.

## Introduction

SARS-CoV2 infections can cause serious diseases in susceptible patients, including elderly and immunosuppressed individuals and subjects with underlying medical conditions. Immunotherapy with selected monoclonal antibodies (mAbs) that possess potent neutralizing activity against the infecting strains has been shown to be an effective treatment for such individuals. For antibody therapies to be useful they must retain potent neutralizing breadth against mutant SARS-CoV2 lineages that emerge in the population. Emergency use authorization was granted by the FDA for several mAb combinations that were highly effective against the initial CoV2 pathogen and its early variants; however, these were ineffective against the Omicron subvariants (Arora et al., 2022; Desautels et al., 2022; Ryuta Uraki, 2022; VanBlargan et al., 2022). More recent mAb therapeutics (Evusheld and Bebtelovimab) were developed that were effective against the initial Omicron isolate, but not against later variants, including the heavily mutated and highly transmissible BQ.1.1 and XBB.1.5 Omicron subvariants and their derivatives that are presently driving CoV2 infections in the US and worldwide (GISAID, 2023; Nowcast, 2023). At the time of writing, XBB.1.5 and its derivatives accounted for > 99% of total infections in the US and >70% of all new infections worldwide. In addition to the loss of sensitivity to mAb immunotherapeutics, these immune-evasive variants were relatively resistant to plasma induced by the original B.1 and BA.5 spike-based bivalent vaccines (Kurhade et al., 2022; Planas et al., 2022; Qu et al., 2022), thus increasing the threat of breakthrough infection and possibly leading to increased mortality in immunocompromised individuals. These findings highlight the need for more effective mAb immunotherapeutics and improved vaccines that can treat and provide sterilizing CoV2 immunity against current and future infections.

A key feature of broadly neutralizing anti-viral antibodies against HIV-1 Env is affinity maturation mediated by the accumulation of somatic hypermutations that improve affinity for a continuously changing antigen (Klein et al., 2013). In contrast to HIV-1 Env, the ancestral CoV2 spike antigen is recognized with high affinity by many germline Ig sequences, and therefore affinity maturation was not required for the generation of potently neutralizing antibodies isolated from early CoV2 infections or from a naïve B cell library (Ferrara et al., 2022; Rogers et al., 2020). However, antibodies isolated from COVID-19 convalescent individuals six months after their PCR diagnosis had more diverse sequences and a greater number of somatic mutations compared to antibodies isolated from convalescent individuals earlier (1.3 months) after their COVID-19 diagnosis (Gaebler et al., 2021), and we and other have shown that the plasma of CoV2-infected vaccinees possessed more potent neutralizing activities against Omicron compared to plasma from uninfected vaccinees or infected-unvaccinated individuals (Choudhary et al., 2022). This suggests that multiple vaccine boosts combined with circulating TfH and de novo T cell responses with CoV2 infection (Koutsakos et al., 2022; Koutsakos et al., 2023), can induce multiple cycles of affinity maturation that lead to the development of hypermutated bnAbs (Goenka et al., 2014). To test this, we screened RBD-binding memory B cells from vaccinated subjects who subsequently recovered from Omicron infections, for the production of broadly neutralizing mAbs against CoV2. This report describes one highly mutated mAb isolated from such a subject that potently neutralizes the Wuhan strain and multiple variant lineages, Including Omicron subvariants extending up to BQ.1.1, a highly immune-evasive subvariant that is still causing a significant percentage of SARS-CoV2 infections worldwide (GISAID, 2023; Wang et al., 2023).

## RESULTS

### The majority of Omicron-neutralizing plasma antibodies target RBD epitopes that are conserved between CoV2-wt and Omicron subvariants

It is known that typical vaccine sera, and sera from unvaccinated subjects infected with early CoV2 variants possessed relatively weak neutralizing activities against the Omicron variants (Cao et al., 2022b), while hybrid immunity induced by breakthrough infection in vaccinated individuals better correlates with effective protection against reinfection and severe disease (Bobrovitz et al., 2023). To investigate the effect of CoV2 infections on the neutralization breadth and potency of vaccinee sera against CoV2-wt (B.1) and Omicron (BA.1) variants viral pseudotype neutralization assays were performed with eight plasmas obtained within 6 months of vaccination from individuals who were subsequently infected, half during the earlier B.1.1.7 and B.1.617.2 waves, and half during the initial Omicron surge. All eight plasmas possessed high neutralization titers against the early B.1 SARS-CoV2 variant, but significantly lower titers for the initial Omicron BA.1 variant. Whereas the decrease in neutralization potencies for the BA.1 vs B.1 virus was >10-fold for the non-Omicron infected vaccinees, a decrease of only 2 to 3-fold was seen for the group infected with Omicron subvariants (**Fig. 1A).** This suggested that Omicron infections produced a greater Omicron-reactive protective antibody response in CoV2 vaccinees than infection with the earlier variants.

**Figure 1:**
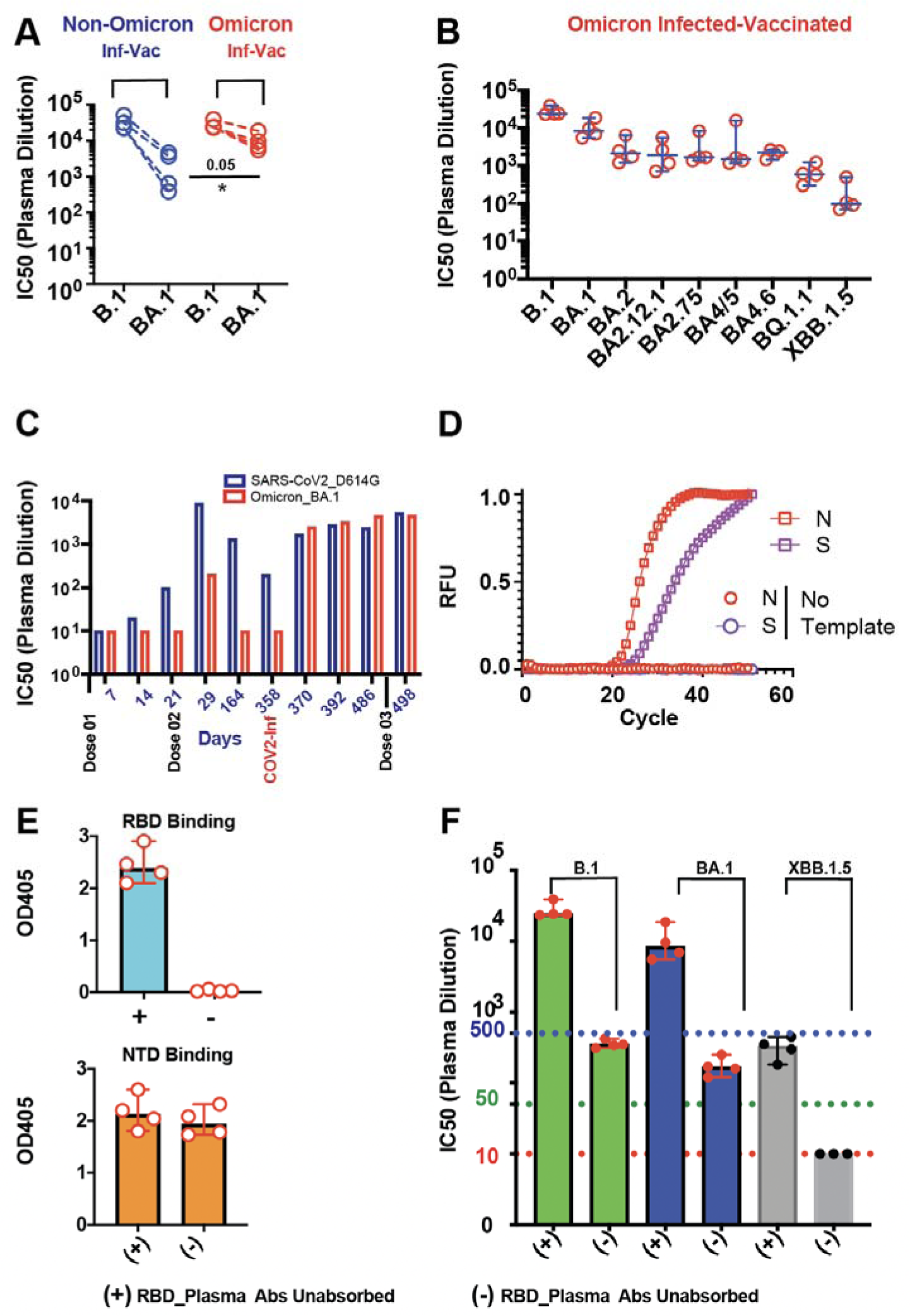
**A**. An assessment of the BA.1 plasma Abs neutralization potency for subjects who have received COVID-19 vaccines and were subsequently infected and recovered from non-omicron SARS-CoV2 infections (blue symbols) and Omicron subvariant infection (red symbols). **B.** A comparison of plasma neutralization potencies of COVID-19 vaccinated subjects recovered from Omicron subvariant infections, against multiple SARS CoV2 variants. **C.** An assessment over time of the effectiveness of the COVID-19 mRNA vaccine in a healthy adult after the first, second, and third doses, as after a subsequent break-through BA.1 infection. **D.** qRT-PCR of the nucleocapsid (N) and Omicron-specific Spike (S) E484A mutation performed on RNA isolated from the nasal swab of a symptomatic CoV2 infected patient, using single nucleotide discriminating becons demonstrating that infection was with the Omicron variant. **E.** A comparison of the RBD-B.1 and NTD-B1 binding in both RBD Abs depleted and undepleted plasma demonstrates the specific depletion of anti-RBD antibodies. **F.** A comparison of the neutralization potency against different CoV2 variants in anti-RBD depleted and undepleted plasma. The green dotted line represents the minimum required protective IC50 of plasma for virus neutralization (IC50 ≥1:50 dilution) (McMahan et al., 2020).

The neutralization potencies of four plasmas from vaccinees with putative Omicron-BA.1 infections were tested against newer, more immune-evasive Omicron variants (**Fig. 1B**). There was a further ≅ 5-fold decrease in neutralization potencies for all of these plasmas against the BA.2.12.1, BA.4/5 and BA.4.6 variants compared to the initial BA.1 Omicron variant, and even greater decreases in neutralization potencies for the more resistant BQ.1.1 and XBB subvariants (≅15 and ≅70-fold respectively less susceptible to neutralization compared to BA.1).

A detailed analysis was performed for one subject who received two doses of the Pfizer vaccine and was infected with the initial Omicron variant one year after vaccination (**Fig. 1C)**. Plasma obtained one month after the second vaccine dose possessed significantly higher neutralizing titers for the vaccine strain than for the Omicron BA.1 variant (IC_50s_ 1:8,761 vs 1:202). By day 164 the neutralizing titer for the SARS-CoV2-wt B.1 virus was further decreased, and the titer against the BA.1 variant was undetectable (1:1,337 vs. <1:10). This subject was infected on day 358 after the initial vaccination; at this time the Omicron BA.1 variant was dominant in the area, and the infecting virus was confirmed to be Omicron by a variant-specific PCR assay using molecular beacon technology as described (**Fig.1D**) (Dikdan et al., 2022). At the time of infection, the neutralizing titer against the B.1 virus had dropped further below 1:200, but by 30 days post-infection the neutralizing titer had risen to ∼1:2,800 for the B.1 virus and slightly higher for the BA.1_Omicron_ variant (∼1:3,200). These titers were retained for the next four months, and a third dose of the parental Pfizer vaccine at that time resulted in a small increase in the titer against the B.1 virus but no detectable effect on the titer against the Omicron variant.

The strain specificity and fraction of the neutralizing activity that was directed against sites in the RBD in this serum were determined by absorption with Sepharose beads containing immobilized RBD_B.1_ recombinant protein. The extent and specificity of depletion of RBD-specific antibodies were confirmed by comparing the binding of absorbed and unabsorbed plasma to recombinant RBD_B.1_ and a control NTD_B.1_ protein by ELISA. All binding activity to the RBD-B.1 protein was depleted, whereas no loss of binding was observed for the NTD antigen, confirming the specific depletion of RBD-directed antibodies (**Fig.1E**). Removal of the RBD antibodies resulted in a >97% decrease in neutralization titer of the plasma against wt CoV2 and Omicron_BA.1_ pseudoviruses and complete deletion against the very resistant Omicron_XBB.1.5_ subvariant (**Fig.1F**). This indicated that the dominant targets of the cross-CoV2 neutralizing antibodies in this Omicron-infected vaccinee were broadly conserved RBD epitopes, with little if any contribution by Omicron-specific epitopes.

### ACOVIMAB is a hypermutated VH1-58/VK3-20 anti-RBD class I mAb isolated from a CoV2 neutralizing B cell culture, that binds to a class I discontinuous epitope in the RBM

Memory B cells with cell-surface IgG that recognized both the SARS-CoV2 B.1 and Omicron BA.1 RBDs were separated from peripheral blood mononuclear cells (PBMCs) isolated from the COVID19-vaccinated BA.1-infected individual described in **Fig. 2A** by sorting with fluorescently labeled wt and Omicron RBD antigens. Single RBD-specific memory B cells were sorted into 96 well culture plates and incubated for two weeks in B cell growth media, to allow cell proliferation and secretion of the Abs into the culture supernatant. Culture supernatant from most of the wells contained non-neutralizing antibodies (NN), three wells weakly neutralized only the BA.1 Omicron subvariant (IC50 < 1:50) but not the SARS-CoV2 wt B.1, and only one out of the 28 RBD-B.1 and BA.1 binding wells possessed potent neutralizing activity against both CoV2-wt (IC50 ≤ 1:172) and BA.1(IC50 ≤ 1:1,589) pseudoviruses. Heavy and light chain variable regions were amplified from this well following standard Ig-PCR conditions as described (Scheid et al., 2009; Tiller et al., 2008), and the respective variable regions were cloned into IgG1 heavy and kappa light chain vectors to generate a functional antibody, called Acovimab.

**Figure 2:**
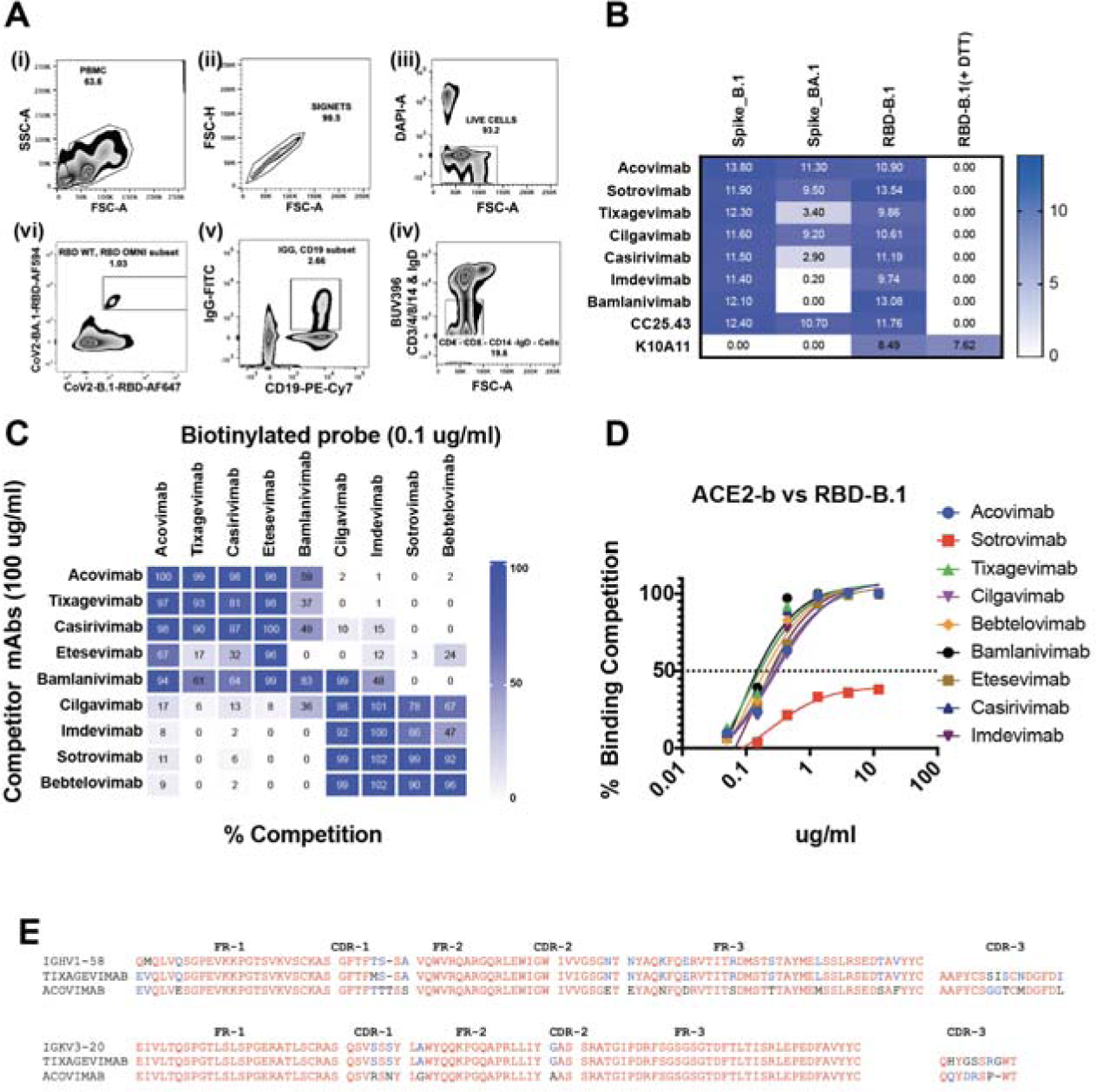
**A**. Schematic presentation of gating strategy to sort RBD-B.1 and RBD-BA.1 bound IgG positive memory B cells. **B.** A comparison of anti-RBD mAbs binding to SARS-CoV2 B.1 and BA.1 spike trimer and determination of linear and conformational epitopes. RBD binding activity is represented by the area under the curve (AUC). **C.** Binding competition assays with Acovimab and eight FDA-approved COVID-19 therapeutic antibodies with RBD-B.1 target. **D.** Binding competition assays with measuring the ability of nine RBD-specific mAbs to inhibit the binding of soluble Hu-ACE2-biotin to RBD-B.1. **E.** Protein sequence alignment of IGHV and IGKV sequences of Acovimab, Tixagevimab and their common germline sequence.

We compared the binding patterns of Acovimab IgG1 to B.1 and BA.1 spike trimers to that of other FDA-approved antibodies and a cross-CoV2 neutralizing monoclonal antibody (CC25.43) provided by the Andrabi lab at Scripps Research. All mAbs tested bound strongly to the trimeric spike and RBD protein of the B.1 variant. Acovimab, Sotrovimab, Cilgavimab and CC25.43 antibodies showed comparably strong binding activity to the BA.1 spike, with much weaker binding signals for Tixagevimab and Casirivimab and no binding signals obtained for Imdevimab and Bamlanavimab against the BA.1 spike **(Fig. 2B).**

Many of the anti-RBD mAbs tested here have been described as having discontinuous epitopes (VanBlargan et al., 2021). To confirm this and to test the Acovimab epitope, we compared the binding of the panel of RBD-specific mAbs to DTT-treated and untreated RBD-gp70 fusion proteins (Datta et al., 2021; Kayman et al., 1999; Kayman et al., 1994), using an anti-gp70 mAb (K10A11) that recognized a linear epitope on the gp70 carrier domain to control for the presence of antigen. None of the RBD-specific mAbs bound to the DTT-treated B.1 RBD-gp70 fusion protein, while the control mAb K10A11 showed comparable binding to both DTT-treated and untreated RBD proteins (**Fig. 2B**). This confirmed that all of the neutralization targets recognized by these antibodies were disulfide bond-dependent conformational epitopes.

Competitive binding assays were then used to define the spatial orientation of the Acovimab target epitope. A total of nine anti-RBD mAbs, including Acovimab and eight well-characterized anti-RBD mAbs that have been previously approved by the FDA for therapeutic use were tested in RBD-B.1 binding competition assays, to identify shared binding competition patterns. Published cryo-EM structural data have shown that the eight therapeutically approved RBD mAbs we tested here fall into three different classes. Tixagevimab, Casirivimab, and Etesevimab are class 1 mAbs, which bind to the RBD on Spike in its up form, Bamlanivimab is a class 2 RBD mAb, which binds to the RBD on the Spike in its down form, while Cilgavimab, Imdevimab, Sotrovimab, and Bebtelovimab are class 3 RBD mAbs, which can bind to the RBD in both it’s up and down forms (Barnes et al., 2020; Bobrovitz et al., 2023; Cathcart et. al., 2022; Desautels et al., 2022; Hansen et al., 2020b; Johnson et al., 2023; Pinto et al., 2020; Zost et al., 2020). These assignments were consistent with the binding competition patterns observed (**Fig.2C**). The three class 1 mAbs cross-competed to the CoV2-B.1 RBD with each other and with Acovimab, showing that Acovimab also belonged to the class 1 group. The four class 3 mAbs also efficiently cross-competed with each other, but not with any of the class 1 mAbs. Bamlanivimab, the only class 2 mAb tested, had a unique competition pattern, as it bridged some of the class 1 and class 3 sites. In addition to competing with all 4 class 1 mAbs, Bamlanivimab also competed with two of the class 3 mAbs (strongly with Cilgavimab and weakly with Imdevimab), but not with either Sotrovimab or Bebtelovimab, the other two class 3 mAbs. Interestingly, competition between Bamlanivimab and Etesevimab was not reciprocal; although Bamlanivimab strongly competed (99% competition) with the binding of Etesevimab, Etesevimab did not block the binding of Bamlanivimab (0% competition). We further confirmed that all of these mAbs except Sotrovimab strongly competed with ACE2 for binding with RBD. As previously reported, Sotrovimab competed with ACE2 very weakly and incompletely **Fig.2D** (Pinto et al., 2020).

Genetic analysis of recombinant Acovimab demonstrated that this antibody was derived from the IGHV1-58 and IGKV3-20 genes; this combination of germline sequences was reported previously for several other CoV2 neutralizing antibodies, including Tixagevimab (AZD8895 or COV2-2196), and BD-836 (Cao et al., 2022b; Dong et al., 2021; Du et al., 2021; Hansen et al., 2020a). Whereas all of the previously described VH1-58/VK3-20 mAbs possessed <5% somatic hypermutations in their IGHV aa sequence, Acovimab IGHV differed from the germline IGVH1-58 sequence by mutation of 15.5% of the aa residues and by the insertion of one aa in VH-CDR1. The Acovimab light chain also possessed a higher level of mutation in its aa sequence (4.5%) compared to the completely unmutated IGKV3-20*01 germline sequence present in Tixagevimab (**Fig. 2E**).

### ACOVIMAB broadly neutralizes CoV2 isolates, including Omicron variants extending to BQ.1.1

Acovimab possessed very strong neutralization activity against the ancestral Wuhan strains and all early CoV2 variants tested (alpha, beta, gamma, delta). In contrast to the previously reported mAbs of the IGVH1-58 class, Acovimab also strongly neutralized many Omicron subvariants (B.1.17, B.1.617.2, B.1.351, P.1, BA.1, BA.2, BA.2.12.1, BA.2.75, BA4/5, BA.4.6 and BQ.1.1), with IC_50s_ of 1-6 ng/ml, and also neutralized the more resistant BA.4/5, BA.4.6 and BQ.1.1 variants, although with lower potencies, with IC_50s_ of 200-653 ng/ml (**Table 1** and **Fig. 3A**). Of note, the neutralization activity of Acovimab for BQ.1.1 contrasted with the complete loss of sensitivity of this variant to Bebtelovimab, which strongly neutralized earlier Omicron variants and had received EUA approval for treatment of CoV2 infected immunocompromised individuals, and to Evusheld (a mixture of Tixagevimab and Cilgavimab), which was approved for pre-exposure prophylaxis of COVID-19 in immunocompromised individuals. However, Acovimab did not neutralize the more recent XBB and XBB.1.5 Omicron subvariants. Comparing the key aa residues between the sensitive and resistant groups of Omicron subvariants reveals that the R346T, N460K and K444T substitutions in the Omicron subvariants had no effect on Acovimab neutralization potency, while the F486V substitution present in BA4/5, BA4.6 and BQ.1.1 resulted in a reduction in potency and the F486S and F486P mutations present in XBB and XBB.1.5 respectively led to a complete loss of binding and neutralizing activity (**Fig. 3A**). A critical role of the F486 residue for class 1 anti-RBD antibody recognition was further demonstrated by the total loss of neutralization potency of Acovimab, Tixagevimab and Casirivimab when the F486A substitution was introduced into the B.1 sequence (**Fig. 3B)**.

**Figure 3:**
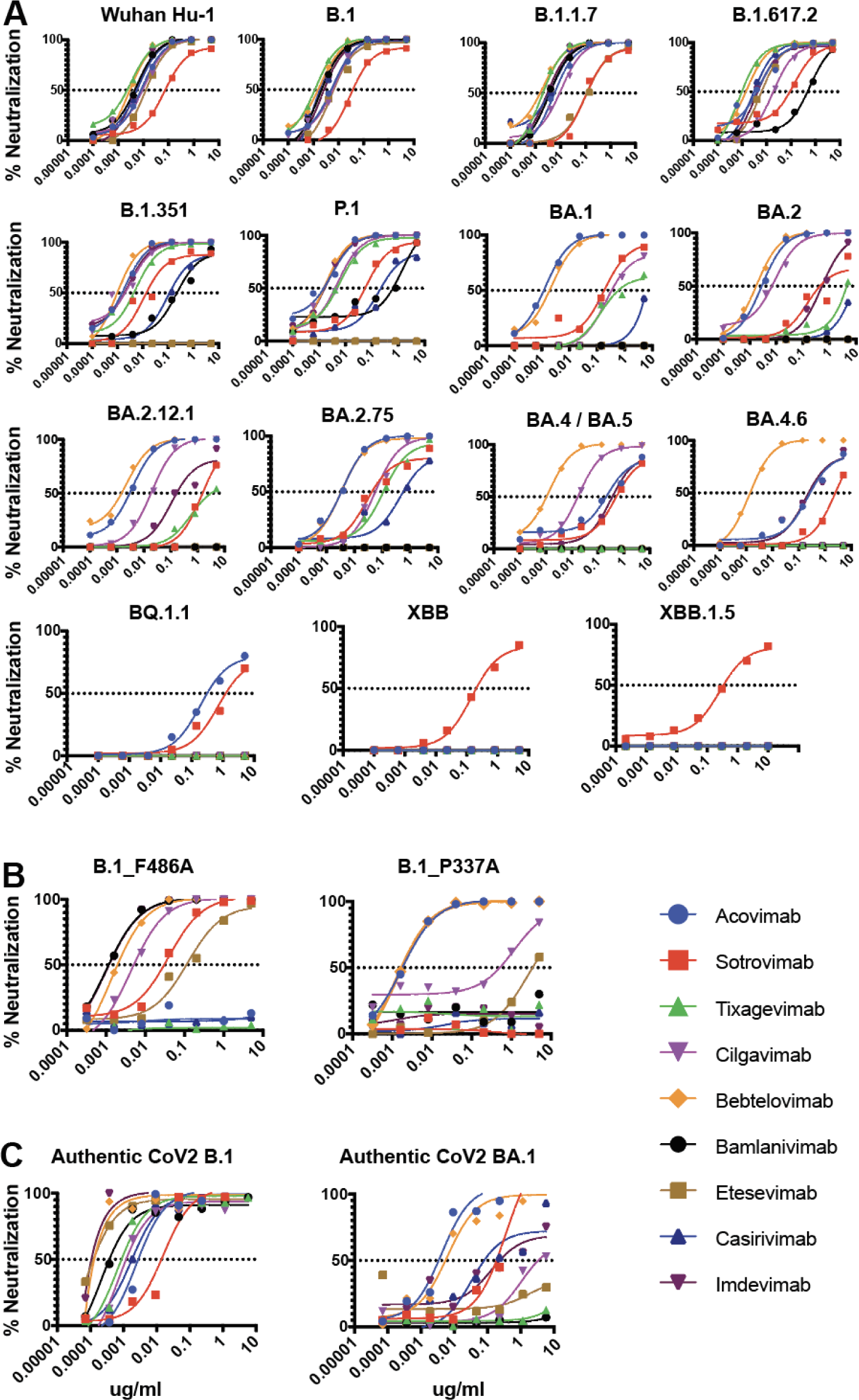
**A**. Neutralization curves of nine anti-RBD mAbs against fifteen SARS-CoV2 pseudoviruses, ranging from B.1 to XBB.1.5. **B.** Neutralization curves of nine anti-RBD mAbs against B.1 with mutant RBDs containing the F486A and P337A point mutations. **C.** Neutralization curve of nine anti-RBD mAbs against B.1 and BA.1 authentic SARS-CoV2 viruses.

**Table 1.**
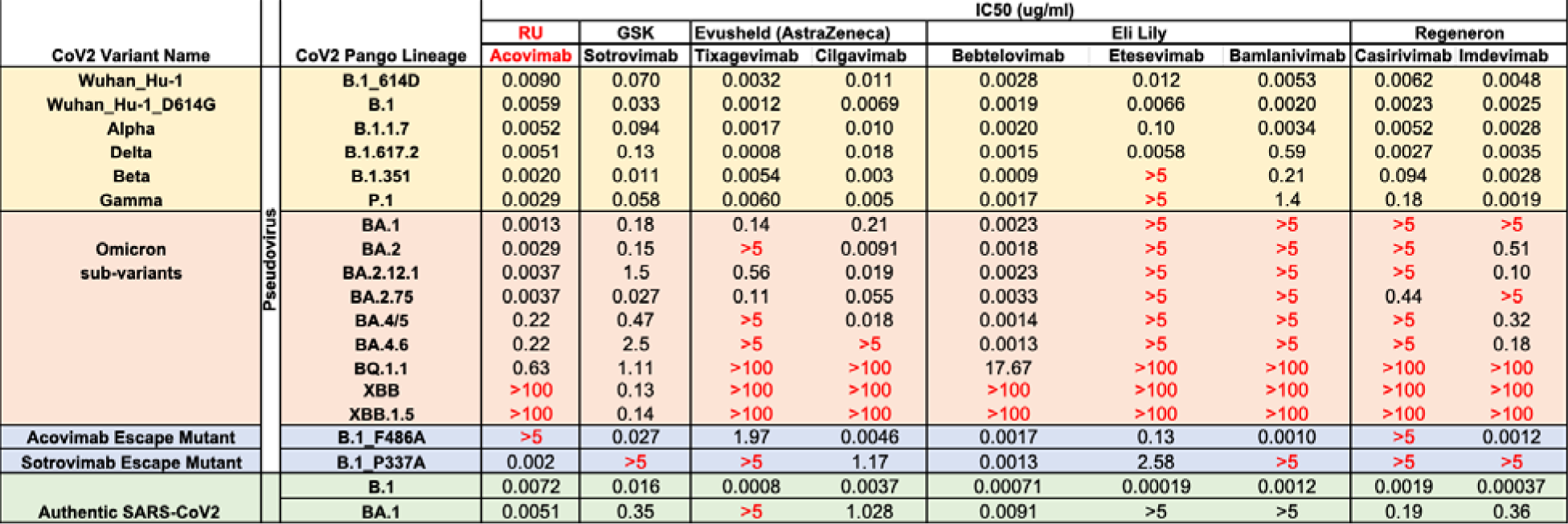
50% Inhibitory concentration (IC50) of nine anti-RBD mAbs against fifteen pseudoviruses ranging from B.1 to XBB.1.5, two B.1 mutants with point mutations in the RBD, and the authentic SARS-CoV2 B.1 and BA.1 viruses.

| CoV2 Variant Name | CoV2 Pango Lineage | IC50 (ug/ml) |  |  |  |  |  |  |  |  |
| --- | --- | --- | --- | --- | --- | --- | --- | --- | --- | --- |
|  |  | RU | GSK | Evusheld (AstraZeneca) |  | Eli Lilly |  |  | Regeneron |  |
|  |  | Acovimab | Sotrovimab | Tixagevimab | Cilgavimab | Bebtelovimab | Etesevimab | Bamlanivimab | Casirivimab | Imdevimab |
| Wuhan_Hu-1 | B.1_614D | 0.0090 | 0.070 | 0.0032 | 0.011 | 0.0028 | 0.012 | 0.0053 | 0.0062 | 0.0048 |
| Wuhan_Hu-1_D614G | B.1 | 0.0059 | 0.033 | 0.0012 | 0.0069 | 0.0019 | 0.0066 | 0.0020 | 0.0023 | 0.0025 |
| Alpha | B.1.1.7 | 0.0052 | 0.094 | 0.0017 | 0.010 | 0.0020 | 0.10 | 0.0034 | 0.0052 | 0.0028 |
| Delta | B.1.617.2 | 0.0051 | 0.13 | 0.0008 | 0.018 | 0.0015 | 0.0058 | 0.59 | 0.0027 | 0.0035 |
| Beta | B.1.351 | 0.0020 | 0.011 | 0.0054 | 0.003 | 0.0009 | >5 | 0.21 | 0.094 | 0.0028 |
| Gamma | P.1 | 0.0029 | 0.058 | 0.0060 | 0.005 | 0.0017 | >5 | 1.4 | 0.18 | 0.0019 |
| Omicron sub-variants | BA.1 | 0.0013 | 0.18 | 0.14 | 0.21 | 0.0023 | >5 | >5 | >5 | >5 |
|  | BA.2 | 0.0029 | 0.15 | >5 | 0.0091 | 0.0018 | >5 | >5 | >5 | 0.51 |
|  | BA.2.12.1 | 0.0037 | 1.5 | 0.56 | 0.019 | 0.0023 | >5 | >5 | >5 | 0.10 |
|  | BA.2.75 | 0.0037 | 0.027 | 0.11 | 0.055 | 0.0033 | >5 | >5 | 0.44 | >5 |
|  | BA.4/5 | 0.22 | 0.47 | >5 | 0.018 | 0.0014 | >5 | >5 | >5 | 0.32 |
|  | BA.4.6 | 0.22 | 2.5 | >5 | >5 | 0.0013 | >5 | >5 | >5 | 0.18 |
|  | BQ.1.1 | 0.63 | 1.11 | >100 | >100 | 17.67 | >100 | >100 | >100 | >100 |
|  | XBB | >100 | 0.13 | >100 | >100 | >100 | >100 | >100 | >100 | >100 |
|  | XBB.1.5 | >100 | 0.14 | >100 | >100 | >100 | >100 | >100 | >100 | >100 |
| Acovimab Escape Mutant | B.1_F486A | >5 | 0.027 | 1.97 | 0.0046 | 0.0017 | 0.13 | 0.0010 | >5 | 0.0012 |
| Sotrovimab Escape Mutant | B.1_P337A | 0.002 | >5 | >5 | 1.17 | 0.0013 | 2.58 | >5 | >5 | >5 |
| Authentic SARS-CoV2 | B.1 | 0.0072 | 0.016 | 0.0008 | 0.0037 | 0.00071 | 0.00019 | 0.0012 | 0.0019 | 0.00037 |
|  | BA.1 | 0.0051 | 0.35 | >5 | 1.028 | 0.0091 | >5 | >5 | 0.19 | 0.36 |

Sotrovimab (derived from parental mAb S309 induced by SARS-CoV1 infection (Cathcart et. al., 2022)) stood out as only antibody which could neutralize all of the CoV2 variants, including XBB.1.5. This was consistent with the fact that Sotrovimab binding epitope does not overlap with the highly mutated RBM region of the Omicron family (Pinto et al., 2020), and none of the Omicron subvariants possessed the known Sotrovimab escape mutations at positions P337 and E340 (Cathcart et. al., 2022). It is noteworthy that the P337A mutation has a wide-reaching effect on the spike configuration, as the neutralization activities of all mAbs except Acovimab and Bebtelovimab were diminished against the B.1_P337A_ mutant (**Fig. 3B**). Different mutations around the crucial binding residues of Sotrovimab had some effect on the neutralization potency of Sotrovimab, resulting in variations of its IC_50_ against different CoV2 subvariants. An interesting observation was that the neutralization potency of Sotrovimab against XBB.1.5 was significantly increased compared to the earlier BA.4.6, BA.2.12.1 and BQ.1.1 variants, by 18-, 11-, and 5-fold respectively (**Table. 1**).

We further examined the neutralizing capacity of four out of eight anti-RBD mAbs against the live B.1 and BA.1 SARS-CoV2 viruses, to verify the correlation between the potencies of RBD mAb-mediated neutralization of spike pseudotyped viruses and the corresponding authentic SARS CoV2 viruses. Similar IC_50s_ values were obtained for the live viruses and the corresponding spike pseudotyped viruses (**Fig 3C and Table.1**).

### Synergistic activity of the combination of Acovimab and Sotrovimab in neutralization of the BQ.1.1 Omicron subvariant

Acovimab and Sotrovimab do not compete with each other for binding to the RBD and have roughly comparable effectiveness in neutralizing the Omicron subvariant BQ.1.1 virus. Acovimab binds to key residues in the RBM and neutralizes the virus by competing with ACE2 for binding to the RBD, while Sotrovimab binds outside of the RBM, and its virus-neutralizing mechanism is independent of ACE2 competition (**Fig.2D and 4A**). Based on these properties we hypothesized that the combination of Acovimab and Sotrovimab might act synergistically to neutralize the highly resistant BQ.1.1 virus. A standard method for examining whether a combination of two therapeutics works synergistically with each other is to calculate their combination index (CI) (Chou and Talalay, 1984; Zhang et al., 2016). This is defined by the following formula: CI (at a given % neutralization) = (D)_1_/(D_X_)_1_ + (D)_2_/(D_X_)_2_ + IZ(D)_1_(D_2_)/(D_X_)_1_/(D_X_)_2_, where (D_1_) and (D_2_) are the concentrations of HuMAbs 1 and 2, respectively, in the mixture used to achieve a given percent neutralization, and (D_X_)_1_ and (D_X_)_2_ are the concentrations of HuMAbs 1 and 2, respectively, used alone to achieve that same percent neutralization. For HuMAbs having different modes of action or acting independently (mutually nonexclusive) IZ=1, whereas for HuMAbs having the same or similar modes of action (mutually exclusive) we assume that IZ=0. CI values are routinely determined using the CompuSyn software program (Chou, 2010), and CI values of less than 1 indicate synergy while values above 1 reflect antagonism.

These assays demonstrated a strong synergism in the neutralization potency of the Acovimab and Sotrovimab combination against the BQ.1.1 pseudovirus, compared to that of the individual antibodies. The calculated neutralization values for Acovimab, Sotrovimab and their combination in a 1:1 ratio are shown in **Fig. 4B and Table. 2**, and the graphs in Fig.4C is showing levels of combination index (CI) and % neutralization at different antibody doses is shown in **Fig. 4C**. These values were calculated with the CompuSyn software program and additional details of this analysis are provided in **Supplement figure 2**.

**Figure 4:**
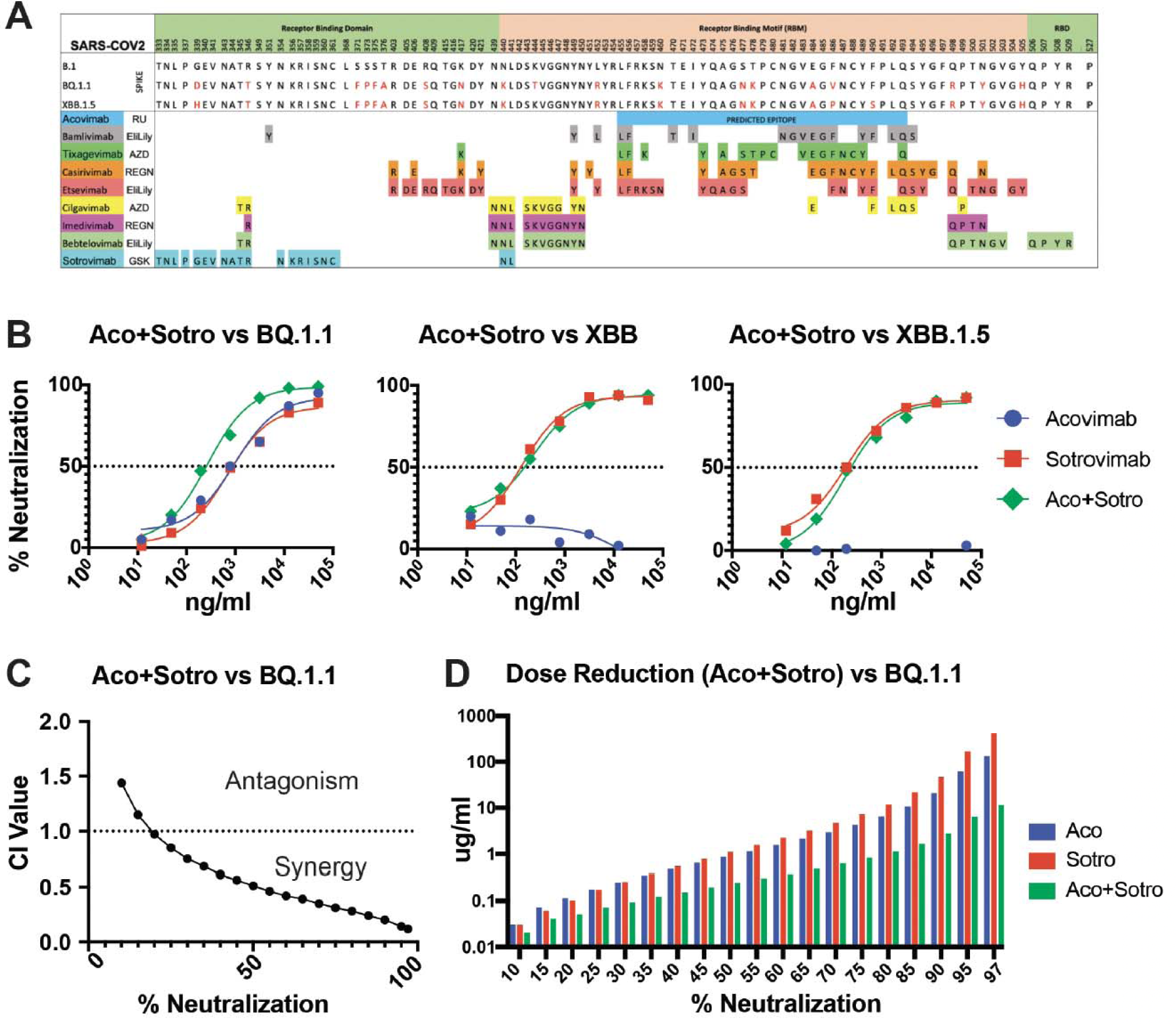
A. Defined epitopes of eight previously described RBD-specific mAbs and the predicted epitope of Acovimab based on its binding competition with the other anti-RBD antibodies. **B.** Neutralization curves of Acovimab, Sotrovimab and their 1:1 combination against BQ.1.1, XBB and XBB.1.5. **C.** Calculated combination index (CI) values plotted for percentage of neutralization, to determine the synergism and antagonism. **D.** Concentration (µg/ml) of the mAbs needed when used alone (red and blue) and in combination (green) to achieve a fixed % neutralization of BQ.1.1. Green bar shows reduced dose needed by the combination of Acovimab and Sotrovimab as compared to their concentration individually required to achieve the same percentage of neutralization of the BQ.1.1 virus.

Our results revealed substantial synergy between Acovimab and Sotrovimab in the neutralization of BQ.1.1, with the strength of synergy increasing at higher levels of neutralization (e.g., CI = 0.97 at 20%, CI = 0.51 at 50%, and CI = 0.11 at 97% neutralization) (**Fig. 4C**). This translated to an impressive reduction in total antibody dose required to achieve near sterilizing levels of protection. When used individually, the concentrations of Acovimab and Sotrovimab required to achieve 97% neutralization (IC_97_ values) were 127 ug/mL and 411 ug/mL respectively, while the same level of neutralization required a combined mAb dose of only 11 ug/mL (**Table 2 and Fig. 4D**), corresponding to a >11-fold dose reduction of Acovimab and >36-fold reduction of Sotrovimab needed to achieve a 97% level of virus neutralization by the combination. This increased potency should dramatically increase the effectiveness of this therapy *in vivo*, and also significantly increase the length of time the treatment would remain effective. As anticipated by the lack of reactivity of Acovimab for these variants, we did not observe any change in the neutralization potency of Sotrovimab against the XBB and XBB.1.5 variants when Acovimab was added (**Fig. 4B)**.

### Potential mechanisms for synergistic neutralization

An essential requirement for the synergistic neutralization effect is that the two mAbs bind to non-competing epitopes. Bebtelovimab weakly neutralizes BQ.1.1, with an IC_50_ of 17.6 ug/ml, and there was no synergistic effect seen when this mAb was combined with Sotrovimab. However, the combination of Bebtelovimab and Acovimab did synergistically neutralize the BQ.1.1 pseudovirus, to a lesser level than that seen for the Acovimab/Sotrovimab combination (**Fig. 5A**). This was consistent with the RBD binding competition results, which demonstrated that Bebtelovimab competed with Sotrovimab but not with Acovimab for RBD binding (**Fig. 2C**). Interestingly, both the Acovimab/Sotrovimab and Acovimab/Bebtelovimab combinations showed modest synergy against the B.617.2 variant (delta), while no synergy was seen for neutralization of the SARS-CoV2 B.1 wt and BA.1 variant with any combinations of RBD mAbs when tested in a 1:1 ratio (**Fig. 5A**).

**Figure 5:**
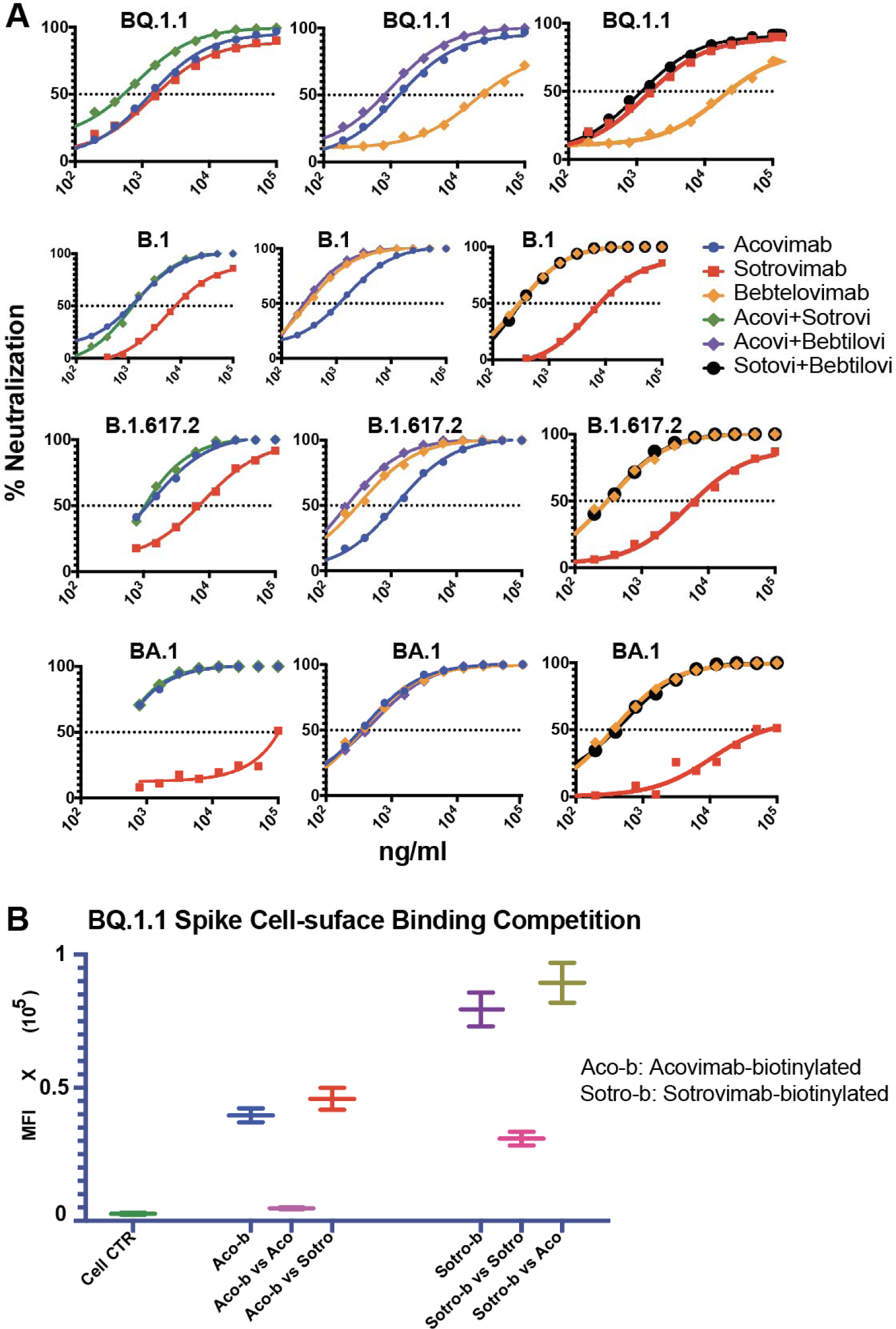
A. Synergistic neutralization by a combination of Acovimab+Strovimab and Acovimab+Bebtelovimab against the BQ.1.1 and B.1.617.2 Delta variant. **B.** Binding competition of biotinylated Acovimab-b and Sotrovimab-b to native BQ.1.1. trimers expressed on the surface of HEK293T cells by 20-fold higher concentrations of unlabeled Acovimab and Sotrovimab.

To understand the mechanism behind the potential synergistic neutralization effect of the Acovimab and Sotrovimab, antibody competitions were performed for the two mAbs against the native cell surface BQ.1.1 Spike trimer. Binding of each biotinylated mAb to the native trimeric spike expressed on the HEK293t cells in the presence of a 20-fold excess of the second unlabeled mAb was quantitated with AF647 labeled streptavidin (**Fig. 5B**). Whereas each of the mAbs efficiently competed with itself, the two mAbs did not cross-compete with each other. This was consistent with the lack of competition of these two mAbs in the RBD binding competition ELISA, and indicated that the two targets were also distant from each other on the native Spike trimer. Interestingly, a low level of enhanced binding was seen for each of the labeled mAbs in the presence of an excess of the second, suggesting that the binding of each of these mAbs had a positive allosteric effect on the exposure of the second site. This further suggested that this enhancement in binding activity may contribute to the synergistic neutralizing activity seen for these antibodies against the BQ.1.1 virus.

### Plasma Abs from persons vaccinated and infected with CoV2 also synergize with Sotrovimab to neutralize the XBB.1.5 subvariant

The neutralizing activities in the plasma of individuals who have recovered from COVID-19 and have been vaccinated are mostly directed towards the RBD (**Fig. 1F**). To more precisely determine the epitope specificity of these neutralizing activities, we examined the ability of the polyclonal plasma antibodies of four vaccinees who were infected with Omicron subvariants to compete for binding of biotin-labeled ACE2 to wt and XBB.1.5 RBD proteins (**Fig. 6A**). Results of these assays showed that three out of four plasma Abs had a comparable ID50 against the binding of ACE2 with RBD-B.1 and RBD-XBB.1.5, while one of the tested plasma competed better for ACE2 binding to the RBD_B1_ than to the RBD_XBB.1.5_ protein. These results suggested that the dominant neutralizing activities of these plasma are aimed at conserved neutralizing epitopes in the RBM that are involved in ACE2 binding.

**Figure 6:**
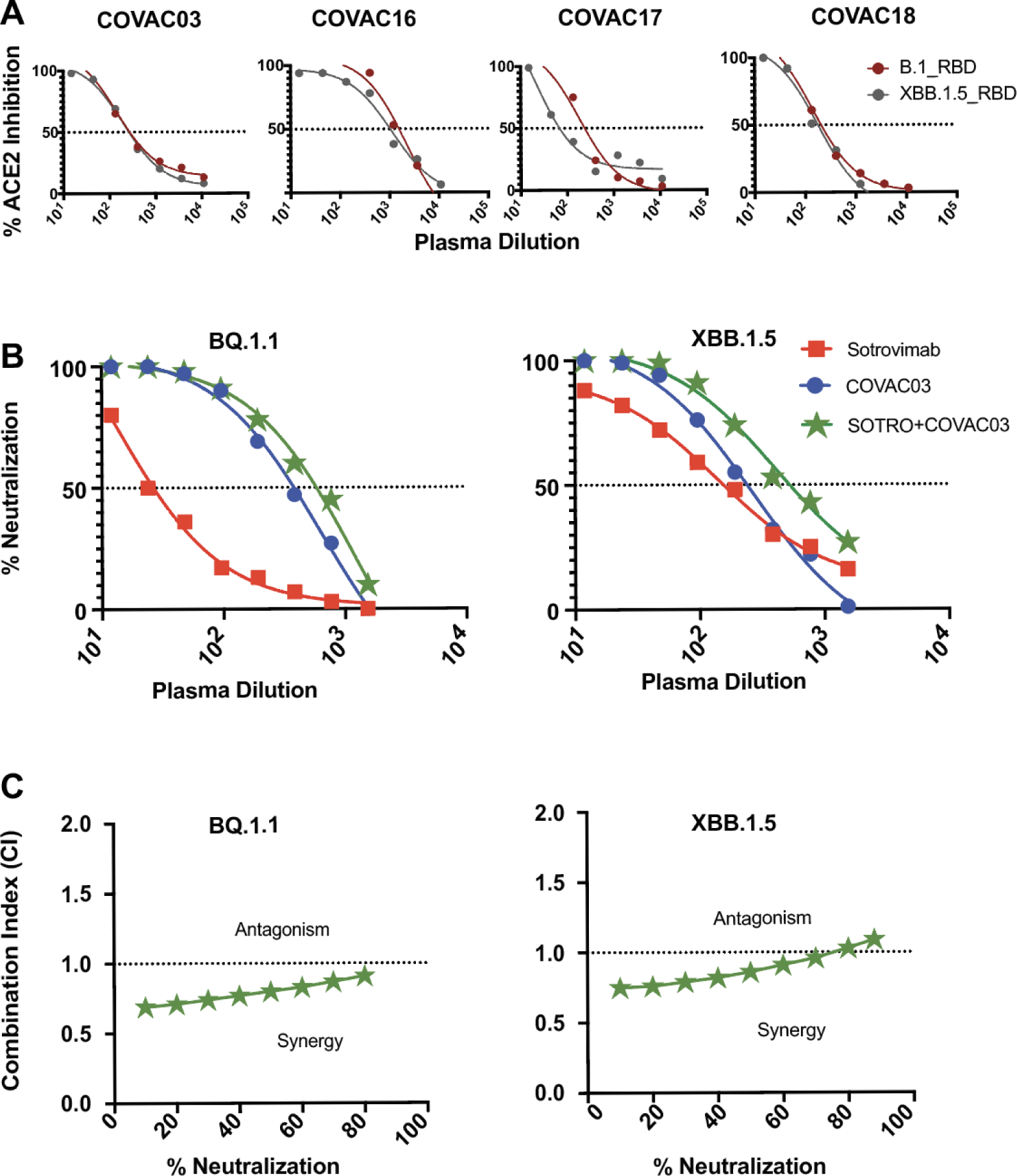
**A**. Binding competition of soluble Human-ACE2-b to RBD-XBB.1.5-gp70 recombinant protein by four polyclonal plasma. **B.** Examining neutralization synergy of COVAC03 plasma Abs combined with Sotrovimab against the SARS-CoV2 BQ.1.1 and XBB.1.5 pseudoviruses. **C.** Combination-Index plot to reflect the synergy or antagonism between the plasma Abs and Sotrovimab against BQ1.1 and XBB1.5

Given that the Acovimab mAb was derived from donor COVAC03 and the plasma Abs responsible for neutralizing BQ.1.1 and XBB.1.5 were mostly directed against the RBM, the ability of the combination COVAC03 polyclonal plasma Abs and Sotrovimab to synergistically neutralize BQ.1.1 and XBB.1.5 was tested. Sotrovimab at 60 ug/ml possessed a similar IC_50s_ as a 1:12 dilution of COVAC03 plasma against the resistant subvariant XBB.1.5 (**Fig. 6B**), and the combination of these antibodies was titered in 2-fold increments for neutralization activity against both the BQ.1.1 and XBB.1.5 subvariants and compared to the Sortovimab and COVAC03 plasma tested individually. The neutralization curve against XBB.1.5 was significantly shifted to the right for the combination (**Fig. 6B**), indicating more potent activity (dilutions for 50% neutralization were 1:504 for the combination, compared to 1:159 for Sotrovimab and 1:208 for the plasma at the same concentrations), resulting in a CI of 0.86 for the combination. Sotrovimab neutralized BQ.1.1 less potently than XBB.1.5, but synergy was also seen for the combination against BQ.1.1, with a CI value of 0.80 at 50% neutralization (**Fig. 6C**). These results indicated that combining Sotrovimab with the endogenous antibodies present in these patient plasma may have some beneficial effect in neutralizing these highly resistant variants.

## Discussion

Key findings of this report are the broadly neutralizing properties of Acovimab, a novel mAb that recognizes a highly conserved site in the RBM, and the potent neutralization synergy of this mAb for some resistant Omicron isolates when combined with Sotrovimab, an older mAb originally derived from a SARS-CoV1 infection that recognizes a highly conserved site located outside of the RBM.

As described in this report, Acovimab is a novel human mAb specific for a conserved RBD epitope that possesses greater neutralization breadth and potency for Omicron variants than earlier mAbs that have received emergency use authorization to treat CoV2 infections. This antibody was isolated from a CoV2-vaccinated individual who was subsequently infected by an early Omicron variant and is derived from the VH1-58/VK3-20 germline genes, which are common precursors for antibodies with potent neutralizing activities that are directed against sites in the RBD ridge (Reincke, Science 2022). In contrast to previous mAbs of this lineage (e.g., Tixagevimab) Acovimab possesses an atypically high level of somatic hypermutation (15.5% for the VH chain and 4.5% for the VL chain); the level of mutation for Tixagevimab is considerably lower (3% for the VH chain and 0% for the VL chain). Both Acovimab and Tixagevimab possess highly potent neutralizing activities for the pre-Omicron isolates and variants, but while Tixagevimab has only weak or no neutralizing activity for the Omicron lineage of variants, Acovimab retains its great potency for the early Omicron BA.1 and BA.2 related sub-variants (IC_50_s of 0.001-0.004 µg/ml), and has lower but still significant neutralization strength (IC_50_s of 0.2-0.6 µg/ml) for a number of the later Omicron sub-variants (BA.4/5, BA.4.6, BQ.1.1).

Residues G485, F486 and N487 in the RBM have been shown to interact with the hydrophobic pocket made by Tixagevimab, and it is reasonable to assume that these positions may also interact with Acovimab. The greater neutralization breadth of Acovimab may be due to its residual affinity to the F486V mutation present in the initial Omicron isolate and its earlier variants. Unfortunately, the activity of Acovimab does not extend to XBB and XBB.1.5, which have acquired different mutations at this position. The XBB variant has the F486S substitution; this negatively impacted affinity for ACE2 (Wang et al., 2023), and this variant quickly evolved by a point mutation into XBB.1.5, which contains the F486P substitution and has comparable ACE2 binding affinity as BA.1 (Cao et al., 2022a; Starr et al., 2020). The dual advantages of high ACE2 affinity and resistance to vaccine-induced immunity allowed XBB.1.5 and its subsequent derivatives to currently dominate CoV2 infections in the US (Nowcast, 2023; Yue et al., 2023).

A particularly interesting finding of this study was the potent level of neutralization synergy seen for the Acovimab/Sotrovimab antibody combination against the highly resistant BQ.1.1 variant, which is circulating in other regions of the world. Sotrovimab (VIR-7831) is an engineered version of human monoclonal antibody S309 (Pinto et al., 2020), and is unique among the reagents that received emergency use approval in that it was generated by a SARS-CoV1 infection and recognizes a highly conserved site outside of the RBM that is retained in the XBB.1.5 variant. Sotrovimab received initial EUA as monotherapy in May 2021, but this authorization was restricted in February 2022 in geographic regions where infections were likely to have been caused by a non-susceptible SARS-CoV2 variant, and completely removed for all US regions in April 2022 due to the spread of resistant Omicron variants, for which Sotrovimab possessed only modest levels of neutralization potency. However, as shown in this report, Sotrovimab actually has a higher neutralization potency for XBB.1.5 (IC_50_ of 0.14 µg/ml), which is similar to its activity against many of the pre-Omicron variants, and better than its activity for the earlier Omicron variants (except for BA.2.75), suggesting that it may be worth reevaluating the utility of this antibody against current infections.

Acovimab and Sotrovimab do not compete for binding to the RBD (**Fig. 2C**), an important requirement for synergy. Interestingly, we revealed a favorable change in the conformation of the native Spike trimer in the presence of these antibodies which leads to modest increases in the binding of the other antibody (**Fig. 6A-B**). The level of synergy seen for neutralization of the BQ.1.1 variant by this antibody combination was particularly impressive at higher levels of neutralization, reducing the total antibody concentration required to achieve 97% neutralization by >11-fold for Acovimab and >36-fold for Sotrovimab (**Supplement Table. 2**). In addition to the beneficial effect of synergy on neutralization potency, the addition of Acovimab should prevent the rapid outgrowth of escape mutants that limited the utility of Sotrovimab monotherapy. Treatment of infections driven by Delta and Omicron variants by Sotrovimab monotherapy resulted in the rapid selection of escape mutants at positions P337 that were completely resistant to this mAb (Rockett et al., 2022; Vellas et al., 2022). These mutations have no effect on the potent activity of Acovimab (**Fig. 3A**), and thus the antibody combination should prevent their generation.

The Omicron isolates in general, and the current XBB.1.5 variant, in particular, are extremely resistant to vaccine-induced antibodies, thus accounting for their frequent reinfections and rapid spread throughout the population. Previous studies have shown that hybrid immunity (vaccination preceded or followed by vaccination) is more effective at inducing antibodies capable of neutralizing Omicron variants (Cao et al., 2022a; Koutsakos et al., 2022), but it was not clear from past studies if this extends to XBB.1.5, and if the residual neutralizing activity is RBD directed. Data presented in this study shows that many of the plasma of infected vaccinees possess lower but significant levels of RBD targeting neutralizing antibodies against XBB.1.5, and that these antibodies also synergize with Sotrovimab for neutralization of XBB.1.5 (**Fig. 4E**). The absorption of these sera with the wt RBD antigen removes this activity (**Fig.1F**), indicating that this activity is mediated by cross-reactive RBD-specific antibodies, perhaps a reflection of the ‘original antigenic sin ‘principle (Zhang et al., 2019). The ability of these sera to effectively compete with the binding of RBD XBB.1.5 to ACE2 (**Fig. 4F**) further shows that at least a fraction of these antibodies are directed against sites in the RBM and suggests that interference with receptor binding is the mechanism of neutralization. The low titer of this activity may reflect a lower concentration of the relevant antibodies, or a lower affinity of these antibodies to the target epitopes, and isolation of such antibodies from these blood samples would allow this to be determined.

The relatively low level of neutralizing activity against XBB.1.5 in these plasma may explain the low level of pressure to induce further escape mutants by mutations in the RBD. Newer variants of XBB.1.5 that are currently spreading (e.g.XBB.1.9.1, XBB.1.16) instead have mutations in targets mediating T cell (N gene) or innate (ORF9b gene) immunity (GISAID, 2023), raising the possibility that the virus is evolving to a form that may also be resistant to other arms of the immune system (Tarke et al., 2021; Wu et al., 2021). A possible outcome of this scenario is that infections with such variants may not be rapidly cleared, and instead lead to a long-term chronic disease. While the acute effects of current strains appear to be milder than earlier variants, the long-term sequela of these infections is not known, and persistent infections may result in additional organs being affected and lead to additional pathologies. This scenario suggests that better antibody-based immunotherapeutics are still useful and needed. Antibodies with higher potencies against Sotrovimab-related epitopes may be useful therapeutics, and B cells from patients with hybrid immunity may be good sources for RBM-specific mAbs with broader potencies that extend to XBB.1.5 and its derivatives. Appropriate combinations of such mAbs may be useful clinical reagents for the treatment of current and future variants of CoV2 and its permutations.

## SUPPLEMENTAL INFORMATION

All parameters are calculated from the mass action law IZ_a_/IZ_u_ = (D/D_m_) ^m^ or D= D_m_ [IZ_a_/(1-IZ_u_)]^1/m^ (Chou Equation) log (IZ_a_/IZ_u_) = m log (D) -m log (D_m_)

Thus, the Median-effect Plot (MEP): x = log (D) y = log(IZ_a_/IZ_u_), gives the slope m, and the x-intercept logD_m_, then the antilog of X-intercept gives the D_m_ value.

Based on the Combination Index Theorem (CIT) and Median-Effect Equation and Plot, when the combination, when the combination (D)_1,2_ for (D)_1_ and (D)_2_ is P/Q, we got CI = (D)_1_/(D_x_)_1_+(D)_2_/(D_x_)_2_ = (D)_1,2_ [P/(P+Q)]/ /(D_m_)_1_[f_a_/(1-f_a_)]^1/m2^

Therefore, substituting, the m and D_m_ parameters, combination ratio P/Q into the corresponding equation given above, and setting f_a_ = 0.01-0.99, the CI values at all effects levels can be simulated as Fa-CI table or Fa-CI Plot. The default setting for the CompuSyn is f_a_ = 0.05, 0.1, 0.15….0.95 and 0.97.

Based on the dose-reduction index (DRI) equations:

(DRI)1 = (D_x_)_1_/(D_1_), (DRI)2 = (D_x_)_2_/(D_2_)

(DRI)1 = (D_m_)_1_/[f_a_/(1-f_a_)]^1/m^_1_ / (D)_1_, (DRI)2 = (D_m_)_2_/[f_a_/(1-f_a_)]^1/m^_2_ / (D)_2_

Similarly, (DRI)_1_ and (DRI)_2_ values at a particular combination data point can be determined, or at different f_a_ value can be simulated.

Median-Effect Plot (MEP) which is also called Chou-Plot gives better visibility as compared to the Dose-Effect Curve because of linearization (**Suppling 2A-B**). The x-intercept (log Dm) in MEP signifies the potency (the antilog gives the Dm value).

CI<1, =1 and >1 indicates synergism, additive effect, and antagonism, respectively.

The Fa-CI table with Fa increment of 5%; at Fa >20% showed a synergistic effect (CI<1).

For BQ.1.1 neutralization, synergism (CI<1) at high dose (higher % neutralization) is more relevant as therapeutic antibodies combination than the CI values at a low dose (low effect).

DRI >1 and <1 indicate favorable and not favorable dose-reduction; DRI=1 indicates no dose-reduction. At 50% neutralization of BQ.1.1, it requires 0.83 ug/ml of Acovimab, and requires 1.08 ug/ml of Sotrovimab. However, it requires 3.48-fold less Acovimab plus 4.52-fold Sotrovimab to achieve the same 50% neutralization of the BQ.1.1 virus (**Suppl. Table 2**).

## ACKNOWLEDGMENTS

We would like to thank members of the Flow Cytometry and Immunology Core Laboratory at NJMS, in particular: Sukhwinder Singh and Tammy Mui-Galenkamp for assistance with FACS sorting. We also thank Erika Kalu at Medical City Dallas Hospital for arranging the timely shipment of blood samples and providing the clinical details. This work was supported by funding from the Center for COVID-19 Response and Pandemic Preparedness (CCRP2), Rutgers University. Graphical abstract was designed by Alok Choudhary.

## AUTHOR CONTRIBUTIONS

A.C. and A.P. planned and designed the experiments. A.C. performed RBD bound memory B cell sorting, in-vitro proliferation, mAb cloning, neutralization, FACS and Data analysis. D.C. made the Spike mutants and mapped the residue sensitive for Acovimab binding and helped with cell-surface Spike trimer/anti-RBD-mAb binding assay. W.H. made the pseudovirus stocks, performed the RBD mAb and ACE2 competition assays, the RBD depletion assays of plasma, and assisted with the RBD mAb competition assay against the BQ.1.1 Spike trimer. A.K. performed the authentic virus neutralization assay. R.J.D. provided help with single nucleotide discriminating molecular beacon PCR. D.J.,G.S. and T.C. helped with the mutagenesis experiment to produce Spike plasmids representing different SARS-CoV2 variants. C.S.,R.J.D., and V.A. helped with blood processing, PBMC isolation, binding ELISA with plasma, and plasmid DNA preparation. A.M., A.R., S.K., M.L., and A.N., provided the FDA-approved therapeutic mAbs and blood samples from vaccinated COVID-19 convalescent individuals. S.S. supervised the SARS-CoV2 neutralization assays. A.C. R.A. and D.B. supervised the CoV2 variants spike production. T.C. analyzed the data for synergistic neutralization by a combination of RBD mAbs. A.C. and A.P. interpreted data, conceived, and supervised the studies. A.C. wrote the paper; A.P. edited the paper; and all authors reviewed the draft.

## STAR METHODS

### Antigens used

Recombinant RBD and NTD recombinant proteins representing different CoV2 variants were expressed as fusion proteins with the gp70 carrier domain as previously described (Datta et al., 2021). In brief, a gene fragment of the CoV2-Spike gene encoding the RBD and NTD was synthesized commercially (Integrated DNA Technologies, Coralville, IA) and cloned at the 3’ end of a gene expressing the N-terminal fragment of the Friend ectotropic MuLV (Fr-MuLV) surface protein (SU) gp70 gene in the expression vector pcDNA3.4 (Addgene, Watertown, MA). The resulting plasmid was transfected into 293F cells using the Expi293 Expression System (Thermo Fisher Scientific) according to the manufacturer’s protocol. Commercial preparations of the Omicron Spike (Cat#SPN-C52Hz), and CoV2-D614G Spike (Cat#SPN-C52H9) were purchased from Acro Biosystems.

### Antigen-antibody binding detected by enzyme-linked immunosorbent assay (ELISA)

RBD plasma antibodies were detected using RBD-WT protein-binding ELISA, and recombinant RBD, S1, and spike proteins from CoV2-WT and Omicron BA.1 variants were used to detect memory B cell culture supernatants and for mAb binding studies. In brief, different proteins were coated overnight at 4°C at a concentration of 100 ng/well in 50 µl of bicarbonate buffer (pH=9.8) using U-shaped medium binding 96-well ELISA plates (Greiner Bio-One; Cat#:650001). Non-fat dry milk (2 %) was used to block the ELISA plate and prepare plasma, culture supernatant, and mAb dilutions. Serially diluted plasma, culture supernatant, and mAbs were incubated at 50 µl/well in duplicates for one hour at 37°C in a blocked plate after washing with wash buffer (1XPBS+0.1% Tween 20) and the antigen-antibody binding was detected by Alkaline phosphatase-conjugated goat anti-human IgG detector antibodies (Jackson ImmunoResearch Laboratories, West Grove, Pa) using 50 µl/well diethanolamine (DAE) buffer. **CoV2 pseudovirus (psV) preparation.**

Codon-optimized D614G, alpha, beta, gamma, delta, kappa, lambda, and omicron spike gene sequences with 18 aa C-terminal truncations were cloned into the pcDNA3.1 mammalian expression vector. Additional Omicron variant spike protein-expressing plasmids were obtained from the Burton lab at the Scripps Institute. Different CoV2 pseudovirus variants were generated by co-transfecting spikes and pNL4.3. Luc. r^-^e^-^plasmids into 293TN cells to produce the CoV2 psV as described earlier (Ferrara et al., 2022). HuACE2-HeLa cells were infected with the CoV2 psV in DMEM media supplemented with polybrene (10 µg/ml), and infectivity was determined by measuring luciferase activity at 72 h post-infection in cell lysates using luciferase substrate (Britelite, PE). To measure the neutralization potency of plasma, psV dilutions yielding ∼ 100,000 RLU were used to infect Hu-ACE2-HeLa cells in the presence of titrated plasma (**Fig. 1A-C and F).**

### Preparation of CNBr-Sepharose bead columns with immobilized RBD for the absorption of plasma RBD antibodies

Cyanogen bromide (CNBr)-activated Sepharose 4 B(Pharmacia#17-0430-01) beads were hydrated with 1mM HCl for 30 min at room temperature, washed with excess 1X PBS, and incubated overnight with the RBD protein in 1X PBS at 4°C on a vertical rotator. After washing with excess 1X PBS, the RBD-conjugated beads were blocked with 1M ethanolamine (pH 8.0) for two hours at room temperature on a vertical rotator, after which the beads were stored in a refrigerator at 4 °C for plasma RBD antibody absorption experiments. For absorption, 1.0 ml of 10-fold diluted plasma from selected patients was incubated with the beads in a tube overnight at 4°C on a vertical rotator, and the beads were spun down at 300 x g to collect the supernatant. The same dilutions of absorbed and unabsorbed plasma were tested for RBD-WT binding by ELISA to determine the percentage of absorption, and antibody titers against an N-terminal domain (NTD) were used to measure the extent of nonspecific losses in plasma antibody concentrations (**Fig. 1E**). RBD-absorbed and-unabsorbed plasma were stored at −80°C and used for the neutralization experiments. Subjects with a high RBD-specific plasma antibody neutralization breadth and potency against CoV2 variants were selected for isolation of broadly neutralizing anti-RBD antibodies.

### RBD^WT+^ and RBD^BA.1+^ bound memory B cell sorting

Two vials of cryopreserved PBMCs with cell counts of 5 million/vial were recovered from the liquid N2 storage, thawed at 37^0^C, and washed with 10% AMEM cell culture medium. Cells were treated with the Fc-block for 5 min at room temperature (RT), followed by incubation at 4^0^C for 30 min with BUV395 tagged mouse anti-human CD3**^+^**, CD4**^+^**, CD8**^+^**, and CD14**^+^** antibodies. After 30 min of incubation, CoV2**^WT^**RBD**^AF647^**, Omicron-BA.1RBD**^AF592^**, anti-human CD19**^PE/Cy7^** and IgG**^FITC^** antibodies of mouse origin were added to the cells, and the tubes were incubated for an additional 30 min in a cold room over the nutator. The total volume of the cell suspension was maintained at <500 µl. After a second incubation at 4 °C, the cells were washed three times with 1XPBS supplemented with 2% FBS and resuspended in 500 µL of 2% AMEM media, and CD19**^PE/Cy7+^**IgG**^FITC+^**CoV2**^WT^**RBD**^AF647+^**BA.1-RBD**^AF592+^** single memory B cells were sorted into flat-bottom culture 96-well plates pre-plated with MS40L cells for in vitro proliferation in the presence of IL2, IL4, IL10, IL21 and CpG (**Fig. 2A**).

### Generation of recombinant broadly CoV2 neutralizing monoclonal antibodies

Cells from wells with anti-RBD antibody binding and neutralizing activity were lysed to prepare cDNA, following the protocol provided by the Invitrogen SuperScript IV First-Strand Synthesis System. Immunoglobulin heavy chain (IGVH) and immunoglobulin light chain (IGVL) were amplified using oligos described earlier (Scheid et al., 2011), and the antibody genes were cloned into the respective human IgG1 heavy and kappa light chain expressing vectors to produce functional human anti-CoV2 mAb.

### Anti-RBD mAb binding competition to RBD-B.1

To identify overlapping epitopes of RBD targeting antibodies, we performed mAbs competition assays with RBD-B.1 (SARS-CoV2-WT). Labeled mAb probes were generated by biotinylation using a standard protocol. In brief, 100 ug of mAb was diluted in 90ul 0.1M bicarbonate buffer (pH 9.6) and 10 ug of biotin (10 mg/ml in DMSO) N-hydroxysuccinimiso-biotin (Sigma # H-1759), mixed well and incubated for 4 hours at RT followed by O/N at 4^0^ C. 10 ul 1M glycine was then added to block any free biotin and incubated for a minimum of 30 min at RT before use. Optionally spin columns were used to remove the free biotin. For the RBD binding competition assays each biotinylated mAbs (100 ng/ml) were competed by unlabeled anti-RBD-mAbs, including an unbiotinylated version of itself, at 100 µg/ml) and % inhibition was determined. The data was plotted as a heat map to resolve major groups based on their binding competition pattern (**Fig. 2C**).

### Competition of binding of soluble biotin-labeled ACE2 to RBD-B.1 by anti-RBD mAbs

To identify if the anti-RBD-mAbs neutralize the SARS-CoV2 by blocking the RBD-ACE2 binding, we examined the ability of mAbs to compete with biotinylated-ACE2 (Acro BioSystem, Cat# AC2-H82E6) for binding with RBD-B.1 (SARS-CoV2-WT) coated on 96-well ELISA plates. MAbs were titrated in duplicate for activity in competing with the binding of 100 ng/ml biotinylated-ACE2 to RBD-wt, and % inhibition was plotted (**Fig. 2D**).

### CoV2 pseudovirus neutralization potency of anti-RBD mAbs

CoV2-psV (pseudovirions) were generated by transient transfection of 293TN cells with codon-optimized spike gene sequences containing 18 aa C-terminal truncations, cloned into the pCDNA3.1 vector. Six earlier CoV2 variants (Wuhan-Hu-1, B.1_D614G, B.1.1.7/alpha, B.1.351/ beta, P.1/ gamma, B.617.2/ delta, and eight Omicron subvariant (BA.1, BA.2, BA.2.12.1, BA.2.75, BA.4/5, BA.4.6, BQ.1.1, XBB, and XBB.1.5) spike genes were used to generate retroviral pseudovirus particles by co-transfecting spike and pNL4.3.Luc.r^-^e^-^ plasmids into 293TN cells. HuACE2-HeLa cells were infected with the CoV2 psVs in DMEM media supplemented with polybrene (10 µg/ml), and infectivity was determined by reading luciferase activity at 72 h post-infection after adding luciferase substrate (Britelite, PE) to the cell lysates. To determine the neutralization potency of mAbs, psV dilutions, producing ∼100,000 RLU were used to infect Hu-ACE2-HeLA cells in the presence of titrated antibodies. The neutralization potency of mAbs, defined as the mAb concentration required to reduce viral infection by 50% (IC50), was calculated using One-Site Fit LogIC_50_ regression (**Fig. 3**).

### Authentic SARS-CoV2 Propagation

Vero E6 cells (obtained from ATCC in Manassas, VA, USA) were grown in DMEM (from Sigma-Aldrich) supplemented with L-glutamine and 10% FBS (from Sigma-Aldrich) to 80% confluency in multiple T75 flasks (from Corning Inc., Corning, NY, USA).After 18 hours, the spent media was decanted, and the cells were washed with sterile PBS (pH 7.2). The SARS-CoV2/USA-WA1/2020 (B.1) and BA.1.529 (Omicron) strain (from BEI Resources, Manassas, VA, USA) was obtained as a lysate of infected cells and diluted in DMEM containing 2% FBS to determine the viral titer. About ∼8 × 106 Vero cells in a T75 flask were infected with 1 mL of the virus suspension and incubated at 37 °C for 1 h, followed by replenishing the cells with 10 mL of DMEM containing 2% FBS. The cell culture supernatant containing the virus was harvested at 72 h post-infection by centrifugation, followed by filtration using a 0.4-micron filter (from Millipore-Sigma, Burlington, MA, USA). Aliquots of the virus-containing media (inoculum) were stored at −80 °C until ready to use and the infectious virus particles in the inoculum were quantitated by plaque assay.

### Authentic SARS-CoV2 Infectivity Assay

The infectivity and titer of the virus inoculum were quantified using a plaque assay with Vero E6 cells. First, 4 × 105 Vero cells were seeded into a six-well cell culture plate in DMEM supplemented with L-glutamine and 10% FBS. After 18 hours, the cells were washed with sterile PBS (pH 7.2) and 400 µL of 10-fold dilutions of the virus inoculum, prepared in serum-free DMEM, was added to each well and incubated at 37 °C with gentle rocking of plates every 15 min for 1 h. The virus inoculum was then removed, and the infected cells were overlayed with 4 mL/well of 1.6% agarose prepared in DMEM with 4% FBS. The plates were allowed to solidify at room temperature for ∼15 min and transferred to a 37 °C incubator with 5% CO2. At 3 days post-infection, the plates were fixed with 10% buffered formalin for 30 min and washed with sterile PBS (pH 7.2). The agar plugs were removed, and the cells were stained with 0.2% crystal violet in 20% ethanol for 10 min. The wells were washed with sterile water and dried, and the clear plaques were counted and presented as the number of plaque-forming units (PFU) of the virus per gram or ml of tissue or lysate.

### Authentic SARS-CoV2 virus Neutralization Assay

To assess the potency of antibodies against the SARS-CoV2 replicative virus USA-WA1/2020 (B.1) and omicron variant (BA1.1.529), a luminescence based method was used to measure the ATP in the viable cells in the presence of virus and antibodies as described earlier (Campbell et al., 2023; Ramasamy et al., 2022; Ravichandran et al., 2020).In brief, serial dilutions of antibodies were incubated in triplicate in 96-well flat-bottom plates with Vero E6 cells and 100 TCID50 of SARS-CoV2 variants in DMEM containing 10% fetal bovine serum. Uninfected cells (Cell Control or CC) and SARS-CoV2 infected cells (Virus Control or VC)without antibody treatment were used as negative and positive control of virus infection. After 72 hours, the plates were equilibrated to room temperature and CellTitre Glo reagent (Cat# G7572, Promega, USA) was added. The luminescence was measured using a Cytation 5 Cell Imaging Multi-Mode Reader. The average luminescence of the antibody titer showing the luminescence values higher than the average of virus control was considered as neutralization and percentage neutralization was calculated with (1-Ab_Lum_ / VC_Avg_)*100 and IC50s was derived using One-Site Fit LogIC_50_ regression.

### Synergy Calculation for Acovimab-Sotrovimab combination against BQ.1.1 neutralization

The neutralization of BQ.1.1 with Acovimab, Sotrovimab, and the combination of the two mAbs in a 1:1 ratio was determined by titrating the antibody 4-fold starting from 50 ug/ml. To assess the allosteric effect of one mAb when the other is already present, 50 ug/ml of Acovimab was added to 50 ug/ml of Sotrovimab. The percentage of neutralization was calculated for each concentration of the mAb, and the data was entered with the assistance of Dr. T.C. Chou, the developer of this method, into the CompySyn software. This is an automated data analysis program based on a mass-action law equations/combination index (CI) algorithm that quantitates synergism or antagonism in drug combinations. The software produces different plots, including the % Neutralization-CI plot (where CI<1 is synergism and CI>1 is antagonism (**Fig. 4C**), and the % Neutralization-DRI (Drug Reduction Index) plot that provides a quantitative value of synergism or antagonism (**Fig. 4D**).

### Selected plasma Abs binding competition with soluble ACE2 to bind with RBD-XBB.1.5

To identify if the anti-RBD plasma Abs that neutralize the SARS-CoV2 XBB.1.5 are RBM or non-RBM directed we performed competition assays with plasma Abs to inhibit the binding of soluble biotinylated-ACE2 to XBB.1.5-RBD-gp70 fusion protein coated on ELISA plates. Four selected plasma from CoV2-Omicron subvariant infected and vaccinated individuals with XBB.1.5 neutralization potency were tested in duplicate by titrations to compete with the binding of 100 ng/ml of biotinylated-ACE2 to XBB.1.5-RBD-gp70 and % inhibition were plotted (**Fig. 5A**).

### Synergy Calculation for Plasma Abs-Sotrovimab combination against XBB.1.5 neutralization

The effect of combining Sotrovimab with plasma Abs that target the XBB.1.5 RBM in a blood donor who had been vaccinated and recovered from the BA.1 Omicron infection was studied. The single donor tested in this experiment was the one whose PBMCs were used to isolate the Acovimab-like mAb. Sotrovimab and plasma dilutions which gave similar IC50s were combined, and serial dilutions were used to measure neutralizing potencies (IC50s) and compared to the activity of similar dilutions of the plasma and mAb by themselves. Neutralization synergies were determined manually using a standard method to calculate their combination index (CI) (Chou and Talalay, 1984; Zhang et al., 2016). This is defined by the following formula: CI (at a given % neutralization) = (D)_1_/(D_X_)_1_ + (D)_2_/(D_X_)_2_ + Ill(D)_1_(D_2_)/(D_X_)_1_/(D_X_)_2_, where (D_1_) and (D_2_) are the concentrations of HuMAbs 1 and 2, respectively, in the mixture used to achieve a given percent neutralization, and (D_X_)_1_ and (D_X_)_2_ are the concentrations of HuMAbs 1 and 2, respectively, used alone to achieve that same percent neutralization. For HuMAbs having different modes of action or acting independently (mutually nonexclusive), Ill = 1, HuMAbs having the same or similar modes of action (mutually exclusive), Ill = 0 (Chou, 2010). CI values of less than 1 indicate synergy for the combination of plasma Abs and Sotrovimab (**Fig. 5C**).

### Binding competition of mAbs to cell-surface expressed trimeric Spike antigens

HEK293T cells pre-plated at 50% confluency in a 100-centimeter cell-culture dish, were transfected with B.1 and BQ.1.1 spike plasmids by mixing the plasmids with TransIT293 transfection reagent (MIR2700) in serum-free media. After 15 hours of transfection, the media was replaced with fresh media, and the cells were harvested after 48 hours of transfection and used for cell-surface spike/mAb binding competition assays. Biotinylated Acovimab and Sotrovimab were used at concentrations of 5 micrograms per milliliter, and their binding to the cells in the presence and absence of unbiotinylated Acovimab and Sotrovimab mAbs (at 100 micrograms per milliliter) was detected with streptavidin-labeled AF647 (SA-AF647) on a benchtop flow cytometer BD-Via. A cell control, which was just treated with SA-AF647, was used to determine the background mean fluorescent intensity (MFI). Data acquired for the mAb binding to the cell-surface trimeric spike were analyzed in Flowjo and the MFI calculated were plotted to evaluate the effect of Acovimab binding to the spike in the presence of excess Sotrovimab mAb, and vice versa (**Fig. 6C**).

### Statistical Analysis

GraphPad Prism (version 8.0) was used to calculate the mean, median, area under the curve (AUC), and to determine suitable parametric or nonparametric tests for statistical analyses of the data. Unpaired Student’s t-tests or Mann-Whitney tests were performed for column statistical analysis to compare two different groups whereas paired t-tests were performed to compare the neutralization potency of plasma against multiple CoV2 variants. The neutralization potency of plasma, defined as the plasma dilution, required to reduce viral infection by 50% (IC50), was calculated using One-Site Fit LogIC_50_ regression. P-values were calculated at a confidence interval of 95% and are indicated as <0.05, or *; <0.01, or **; and <0.001, or ***.

---

**Supplement Figure 1:**
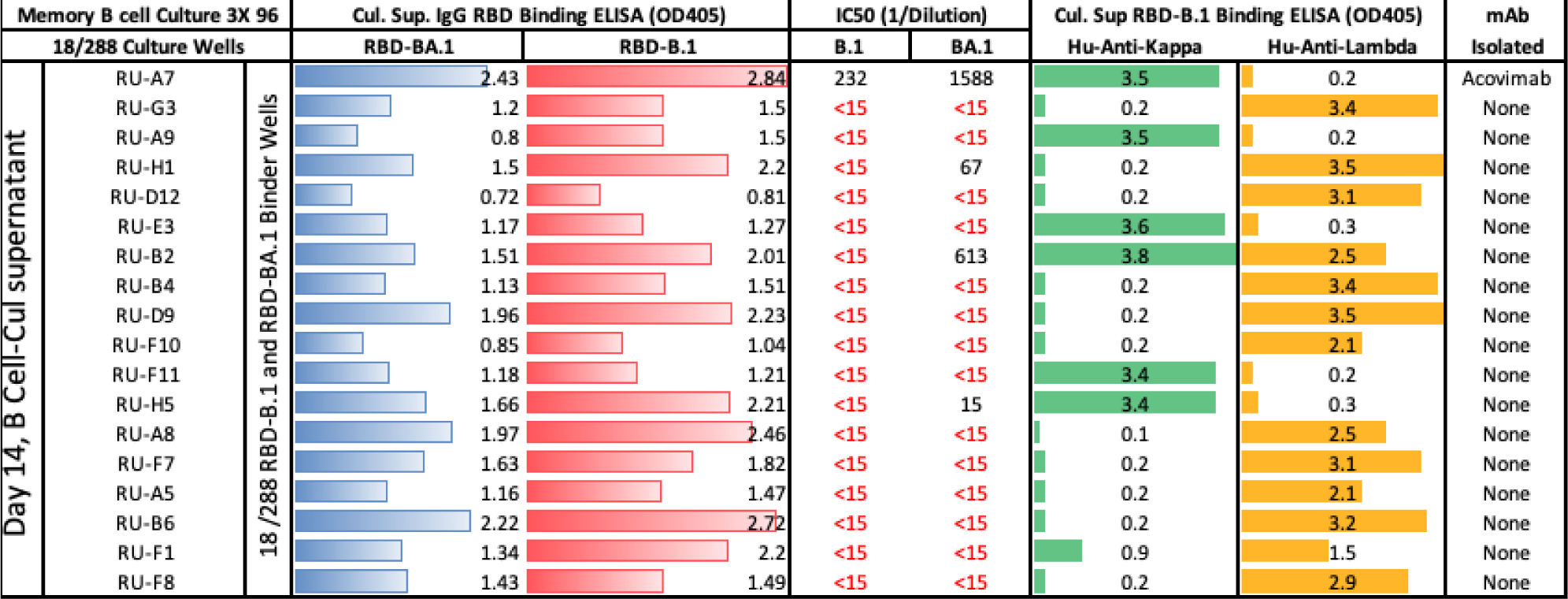
A total of 288 single antigen-bound memory B cells were sorted into three plates pre-plated with MS40L cells in B cell growth media supplemented with cytokines and CpG for in vitro proliferation as described previously (Choudhary et al., 2018). The cells were cultured for two weeks, resulting in approximately 17–18 cycles of cell division (∼ 125,000 cells/well), and the culture supernatants were screened for secreted antibodies binding to RBD-B.1 and RBD-BA.1 by ELISA. A total of 45 RBD IgG-positive wells were detected, and supernatants from 18 bound to both RBD-B.1 and RBD-BA.1 antigens. These wells were further tested for neutralization activity against B.1 and BA.1 pseudoviruses, Acovimab was isolated from well RU-A7; culture supernatant from these cells strongly neutralized the BA.1 viruses, with a 7-fold lower B.1 neutralization titer for BA.1, consistent with the stronger activity of Acovimab for BA.1 over B.1. Weaker neutralizing activities for the BA.1 virus were also found for supernatants from wells RU-H1, RU-H5, and RU-B2, but mAbs were not isolated from these wells. Both Kappa and Lambda RBD-binding Abs were present in well RU-B2, implying that some wells may contain mixtures of RBD-positive B cells which may have contributed to differences in specificities seen in binding and neutralization assays for these wells.

**Supplement Figure 2:**
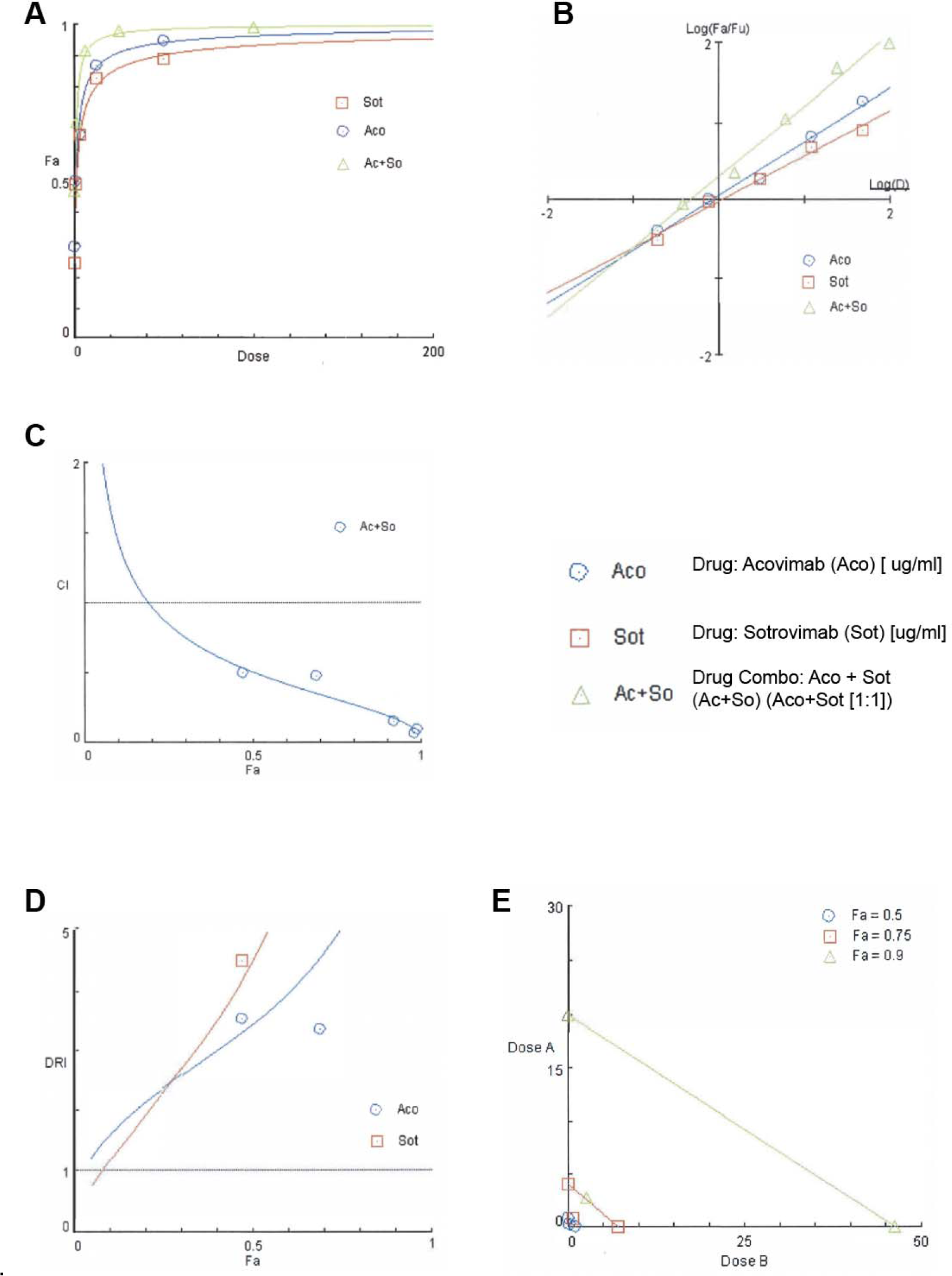
Synergy between two anti-RBD mAbs against BQ.1.1 neutralization was calculated b CompuSyn, a computer program for quantitation of synergism and antagonism in drugs combination. **A.** Dose Effect Curves (DEC) plotted for Acovimab (Blue), Sotrovimab (Red) and their combination in 1:1 (Green) **B.** Median Effect Plot (MEP) plotted for Acovimab (Blue), Sotrovimab (Red) and their combination in 1:1 (Green) **C.** Combination-Index Plot plotted for Acovimab (Blue), Sotrovimab (Red) and their combination in 1:1 (Green), CI<1 = Synergy, CI=1= Additive, CI>1 = Antagonism **D.** Drug Reduction Index (DRI) plotted for Acovimab (Blue), Sotrovimab (Red) and their combination in 1:1 (Green), below zero is non-favorable dose reduction, here both Acovimab and Sotrovimab show favorable dose reduction (>1), which is beneficial as combination therapy. **E.** Isobologram for 50%, 75% and 90% synergism are shown. Combination data point on the diagonal line indicates additive effects, on the lower left indicates synergism, on the upper right indicates antagonism.

**Supplement Table 2:** Fa-CI-DRI Table for Acovimab, Sotrovimab and their combination in 1:1 against BQ.1.1 virus. CI values <1, indicating synergy, start at 20% neutralization (CI = 0.97), with the most significant effects seen for higher levels of neutralization (CI = 0.12 at 97% neutralization).

| Fa | D1 | D2 | D1-2 | Fold Reduction in Combo |  | Comb. Index |
| --- | --- | --- | --- | --- | --- | --- |
| % Neutralization | Acovimab | Sotrovimab | Aco+Sotro | DRI_Aco | DRI_Sotro | CI Value |
| 10 | 0.03 | 0.03 | 0.02 | 1.65 | 1.20 | 1.44 |
| 15 | 0.07 | 0.06 | 0.04 | 1.93 | 1.59 | 1.15 |
| 20 | 0.11 | 0.10 | 0.05 | 2.17 | 1.96 | 0.97 |
| 25 | 0.17 | 0.17 | 0.07 | 2.39 | 2.33 | 0.85 |
| 30 | 0.24 | 0.25 | 0.09 | 2.61 | 2.71 | 0.75 |
| 35 | 0.34 | 0.38 | 0.12 | 2.82 | 3.11 | 0.68 |
| 40 | 0.46 | 0.54 | 0.15 | 3.03 | 3.54 | 0.61 |
| 45 | 0.62 | 0.77 | 0.19 | 3.25 | 4.01 | 0.56 |
| 50 | 0.83 | 1.08 | 0.24 | 3.48 | 4.52 | 0.51 |
| 55 | 1.11 | 1.52 | 0.30 | 3.72 | 5.10 | 0.46 |
| 60 | 1.50 | 2.16 | 0.37 | 3.99 | 5.77 | 0.42 |
| 65 | 2.04 | 3.12 | 0.47 | 4.29 | 6.56 | 0.39 |
| 70 | 2.84 | 4.60 | 0.61 | 4.64 | 7.53 | 0.35 |
| 75 | 4.08 | 7.07 | 0.81 | 5.05 | 8.76 | 0.31 |
| 80 | 6.19 | 11.56 | 1.11 | 5.57 | 10.41 | 0.28 |
| 85 | 10.25 | 20.97 | 1.63 | 6.28 | 12.84 | 0.24 |
| 90 | 20.02 | 46.25 | 2.73 | 7.34 | 16.97 | 0.20 |
| 95 | 59.03 | 165.86 | 6.23 | 9.47 | 26.61 | 0.14 |
| 97 | 127.44 | 411.54 | 11.23 | 11.35 | 36.64 | 0.12 |

**5 data points (average Fa value of triplicate assays) were entered.**
| Data for Drug: Aco [ug/ml] |  | Data for Drug: Sot [ ug/ml] |  | Data for Drug Combo:<br>Aco+Sot [1:1] |  |
| --- | --- | --- | --- | --- | --- |
| Dose | Effect<br>(Fa) | Dose | Effect<br>(Fa) | Dose | Effect<br>(Fa) |
| 50 | 0.95 | 50 | 0.89 | 50 + | 0.99 |
| 12.5 | 0.87 | 12.5 | 0.83 | 12.5 + | 0.98 |
| 3.125 | 0.65 | 3.125 | 0.65 | 3.125 + | 0.92 |
| 0.781 | 0.5 | 0.781 | 0.49 | 0.781 + | 0.69 |
| 0.195 | 0.29 | 0.195 | 0.24 | 0.195 + | 0.47 |
| 5 data points entered. |  | 5 data points entered. |  | 5 data points entered. |  |
| **X-int:** | -0.0797 | **X-int:** | 0.03404 | **X-int:** | -3202 |
| **Y-int:** | 0.05508 +/- 0.04388 | **Y-int:** | -0.0199 +/- 0.03935 | **Y-int:** | 0.28913 +/-<br>0.07580 |
| **m<sub>1</sub>:** | 0.69091 +/- 0.04455 | **m<sub>2</sub>:** | 0.58505 +/- 0.03995 | **m<sub>1-2</sub>:** | 0.90310 +/-<br>0.06501 |
| **Dm<sub>1</sub>:** | 0.83229 | **Dm<sub>2</sub>:** | 1.08154 | **Dm<sub>1-2</sub>:** | 0.47846 |
| **r:** | 0.99382 | **r:** | 0.99308 | **r:** | 0.99232 |

**m:** The slope of the median effect, the dynamic order, or the “shape” of the dose-effect curve; m=1, >1 and <1 indicate the hyperbolic, sigmoidal, and flat sigmoidal shape, respectively.
**Dm:** This is the median effective dose; in this case, it is the IC50, which indicates the neutralization potency of BQ.1.1. The value can be obtained from the X-intercept of the ME-plot.
**r:** The linear correlation coefficient of ME-plot

